# Signaling compartment at the ciliary tip is formed and maintained by intraflagellar transport and functions as sensitive salt detector

**DOI:** 10.1101/861955

**Authors:** Servaas N. van der Burght, Suzanne Rademakers, Jacque-Lynne Johnson, Chunmei Li, Gert-Jan Kremers, Adriaan B. Houtsmuller, Michel R. Leroux, Gert Jansen

## Abstract

Primary cilia are ubiquitous antenna-like organelles that mediate cellular signaling and represent hotspots for human diseases termed ciliopathies. How signaling subcompartments are established within the microtubule-based organelle, and for example support Hedgehog or cGMP signal transduction pathways, remains a central question. Here we show that a *C. elegans* salt-sensing receptor type guanylate cyclase, GCY-22, accumulates at a high concentration within the distal region of the cilium. This receptor uses DAF-25 (Ankmy2 in mammals) to cross the transition zone (TZ) membrane diffusion barrier in the proximal-most region of the ciliary axoneme. Targeting of GCY-22 to the ciliary tip is dynamic, requiring the cargo-mobilizing intraflagellar transport (IFT) system. Disruption of transit across the TZ barrier or IFT trafficking causes GCY-22 protein mislocalization and defects in the formation, maintenance, and function of the ciliary tip compartment required for chemotaxis to low NaCl concentrations. Together, our findings reveal how a previously undescribed cilium tip cGMP signaling compartment is established and contributes to the physiological function of a primary cilium.

## Introduction

The primary cilium is a specialized signaling organelle used by metazoan cells to transduce environmental cues. Signaling proteins important for cilia function must reach and maintain their correct sub-ciliary position and concentration. Failure to do so results in various diseases^1^. Our understanding of the regulation of protein localization in cilia, and their relevance in creating functional signaling domains, remains limited, however.

Cilia use two mechanisms, a trafficking system and a diffusion barrier, that function together to regulate the trafficking of proteins into, within, and out of cilia. The main ciliary trafficking machinery, intraflagellar transport (IFT), facilitates bidirectional transport of cargo, including signaling proteins, from the base/foundation (basal body) to the tip of the axoneme^2^. Anterograde IFT to the tip relies on kinesins, and cytoplasmic dynein enables retrograde transport back^3,4^. Two IFT modules, subcomplexes-A and -B^5,6^, together with another module containing BBS proteins (BBSome) that is thought to bridge the subcomplexes, play essential roles in cargo transport^7,8^. The best-known IFT cargos are axoneme structure components, including tubulin^9,10^, but signaling proteins, like the TRPV channel subunits OSM-9 and OCR-2 in the nematode *C. elegans*, are also transported^11^. Additionally, several mammalian ciliary signaling proteins, namely the GPCR SSTR3 and Hedgehog signaling component SMO, traverse the cilium by both IFT and diffusion^12^.

To help confine proteins to cilia, a subdomain immediately distal to the basal body, called the transition zone (TZ), acts as a diffusion barrier for both membrane and soluble proteins^13–15^. How the TZ acts with IFT or other trafficking systems to regulate the composition of the sensory organelle is not well understood^16,17^.

Signaling proteins can have different sub-ciliary localizations, including the proximal or distal segments, or ciliary tip. For example, the *C. elegans* cyclic nucleotide gated channel TAX-2 localizes to the proximal region adjoining the TZ^18^, while OSM-9 and OCR-2^11^ and several GPCRs^18–20^ localize along the length of the cilium. In mammals, the kinesin-like protein KIF7 and Hedgehog signaling components SUFU and GLI2 localize at the cilium tip^21–23^. Mislocalization of ciliary proteins can impair signaling and development. For example, mislocalization of PDE6 and GRK1 can cause retinitis pigmentosa^24^ and *BBS2* mutant mice display defects presumably caused by mislocalized rhodopsin^25^. Despite their importance, the mechanisms that govern how signaling components concentrate along specific ciliary subdomains remain largely unknown.

To explore the molecular mechanisms underlying ciliary signaling domain formation, maintenance and function, we use the cilium-dependent NaCl response of *C. elegans* as a model system. Its attraction to NaCl is mediated by two bilateral chemosensory ASE head neurons that express receptor-type guanylate cyclases, including GCY-22 (ASER), required for responding to Cl^-^, and GCY-14 (ASEL), for the Na^+^ response^26–29^.

By endogenously tagging GCY-22 with GFP, we discovered that the guanylate cyclase exists at a high concentration in a cilium tip compartment. This localization depends on the IFT machinery, including the BBS complex. We further show that DAF-25 (mammalian Ankmy2 ortholog), is required for GCY-22 ciliary entry. Structure-function studies uncovered GCY-22 protein domains needed for entry and tip localization. Disrupting receptor localization at the tip compartment hinders the ability of *C. elegans* to detect low concentrations of NaCl. Together, our findings provide mechanistic insights into the formation, maintenance and function of a novel ciliary subdomain essential for cGMP-signaling.

## RESULTS

### GCY-22::GFP is highly concentrated at the ASER ciliary tip and periciliary membrane compartment

We used CRISPR/Cas9 to tag the *gcy-22* gene with *GFP* and determine the sub-cellular localization of this receptor in the ASER neuron. Confocal microscopy and co-labeling with markers for the ER (TRAM-1) and trans Golgi (APT-9) revealed that GCY-22::GFP is present within both of these cell body compartments (**Fig. 1a, Supplementary Fig. 1a,b**). In the dendrite, time-lapse imaging revealed anterograde and retrograde transport of GCY-22::GFP (**Fig. 1c**), reminiscent of kinesin and dynein-mediated vesicle transport^30^. Auxin inducible degradation of the dynein heavy chain subunit DHC-1^31^ resulted in stationary particles, showing that GCY-22::GFP dendritic transport is DHC-1-dependent (**Fig. 1c**). At the dendritic terminus, GCY-22::GFP is abundant at the periciliary membrane compartment (PCMC), a vestibule of the cilium (**Fig. 1a**).

**Figure 1.**
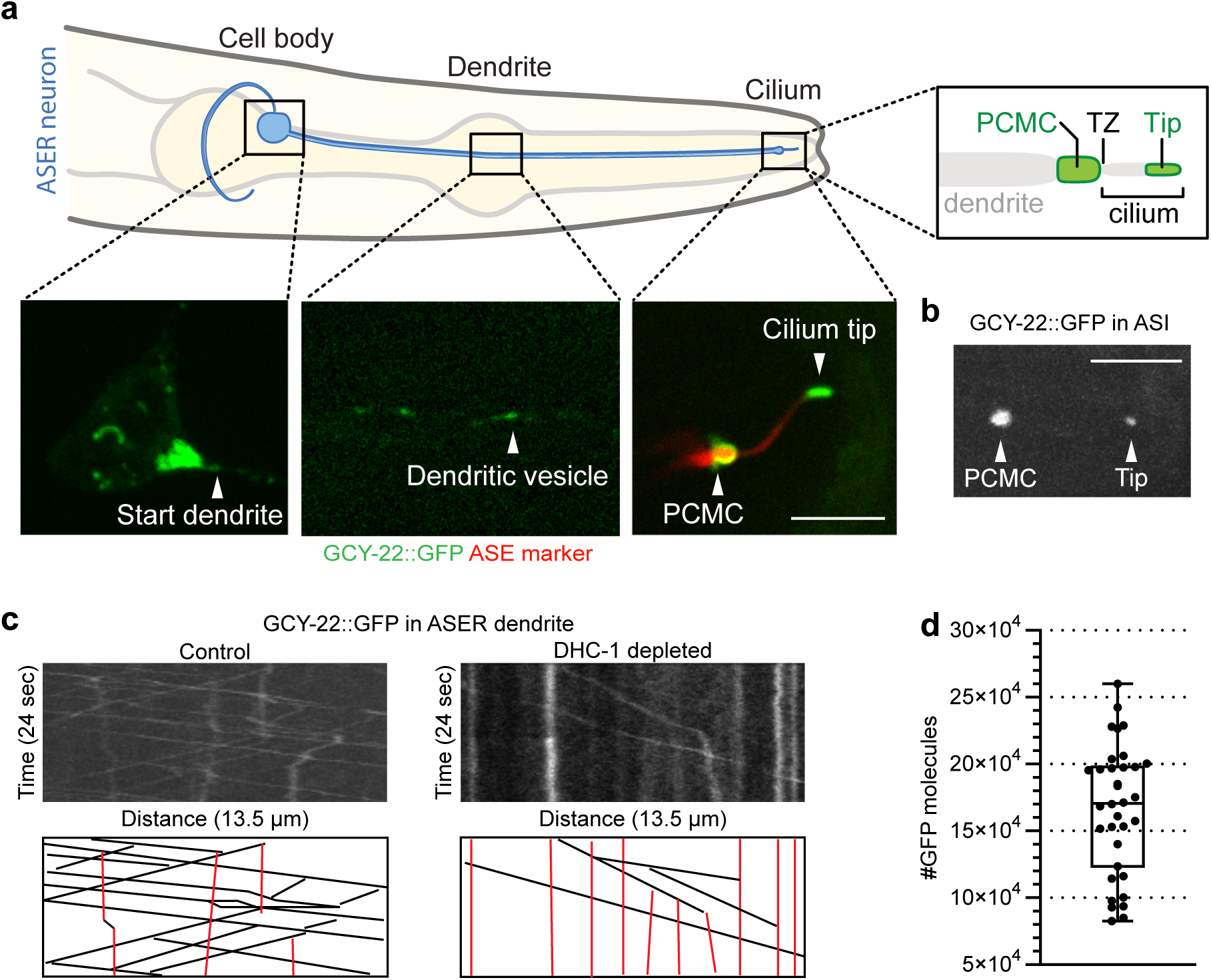
GCY-22::GFP traffics along the dendrite from the cell body to the base of the cilium and concentrates at the ciliary tip in the salt-sensing ASER neuron. **(a)** Schematic of the head of the animal showing the ASER cell body, dendrite and cilium. Inserts: schematic of the cilium depicting the periciliary membrane compartment (PCMC), transition zone (TZ), and cilium tip, and fluorescence images with GCY-22::GFP (green) and mCherry (red) showing localization in the cell body, vesicles in the dendrite and localization to the PCMC and tip of the cilium. **(b)** Ectopic expression of GCY-22::GFP in the ASI neuron showing localization at the PCMC and cilium tip. **(c)** Kymograms showing dendritic transport tracks of GCY-22::GFP in wild-type background and reduced transport in a *dhc-1::GFP::degron* background in the presence of auxin (IAA). Black lines indicate moving and red lines indicate stationary vesicles. **(d)** Quantification of GFP molecules at the cilium tip of ASE neurons. Scale bars indicate 5 μm.

The most striking localization of GCY-22::GFP is its strong enrichment at the ciliary tip **(Fig. 1a**), but not along the axoneme. GCY-22::GFP ectopically expresssed in ASI localized to the PCMC and ciliary tip, although in low amounts (**Fig. 1b**), suggesting a localization mechanism that is not cell specific. Using purified GFP as reference, we estimated the ASE cilium tip concentration of GCY-22::GFP to be 2.77 mM, reflecting an average density of 106,571 (±31,016) molecules/μm^2^ (**Fig. 1d**). Only rhodopsin in mammalian photoreceptor cells has a similar density, of up to 48,300 molecules/μm^2^, which enables photoreceptors to respond to single photons^32–34^. This suggests that the ASE neuron ciliary tip compartment could function as a highly sensitive salt detector.

### GCY-22::GFP exists in stable pools at the periciliary membrane and ciliary tip

To determine how GCY-22::GFP pools at the PCMC and ciliary tip are replenished, we used Fluorescent Recovery after Photobleaching (FRAP). Recovery of the entire PCMC (28% after 25 minutes) was modest, potentially reflecting continuous transport of GCY-22::GFP from the cell body to the dendritic terminus (**Fig. 2a**). Recovery of the entire ciliary tip was slower (7% after 25 minutes), indicating little or slow transport towards the tip (**Fig. 2b**).

**Figure 2.**
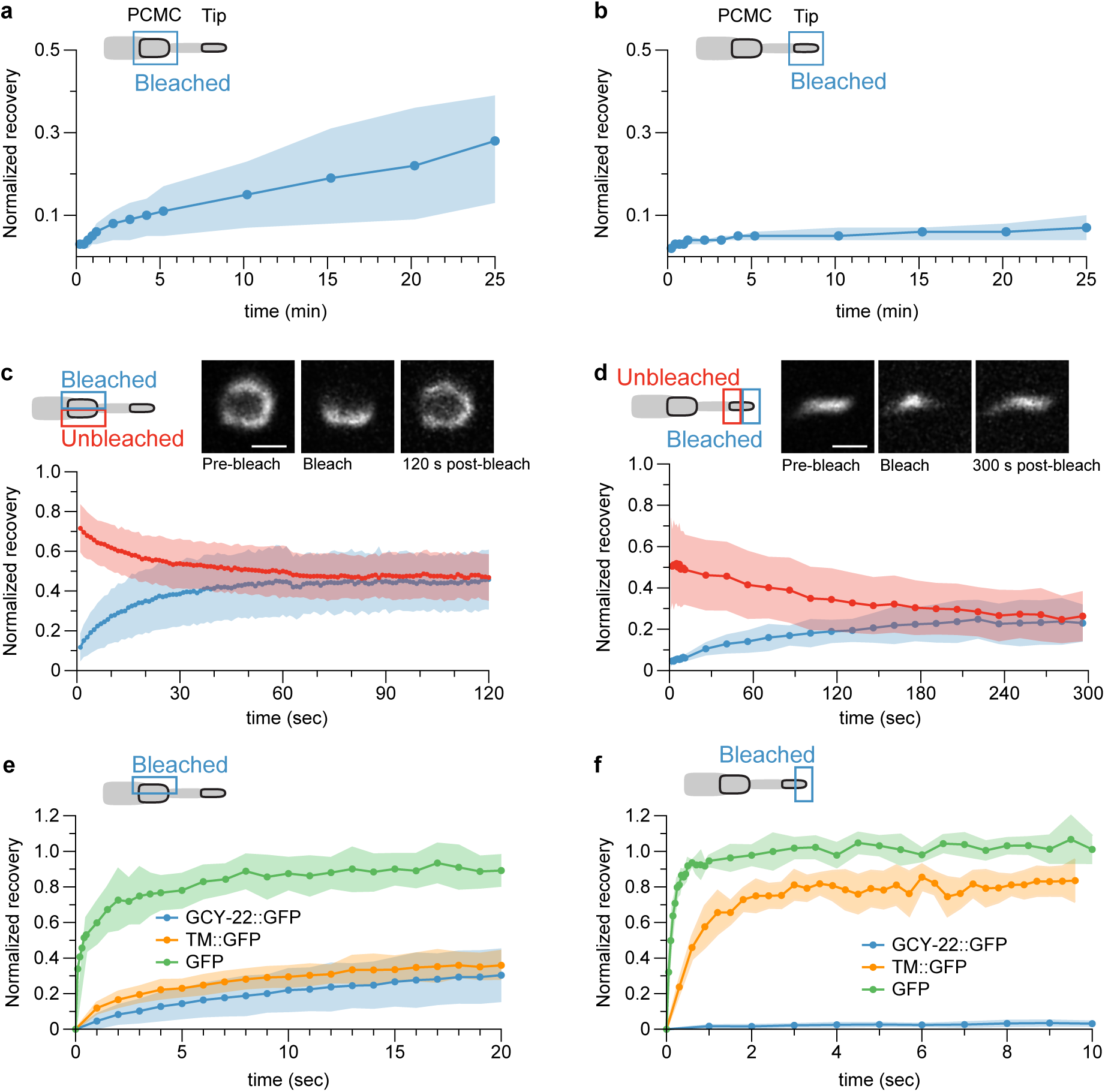
GCY-22::GFP is present in stable pools at the PCMC and cilium tip. **(a)** Fluorescence recovery after photobleaching (FRAP) of the entire PCMC (n=7). **(b)** FRAP of the entire cilium tip compartment (n=8). **(c)** FRAP of half of the PCMC (n=19). Insert: Fluorescence images of the PCMC pre- and post-bleach. **(d)** FRAP of half of the tip compartment (n=6). Insert: fluorescence images of the tip pre- and post-bleach. **(e)** Fluorescence recovery of GCY-22::GFP (blue, n=19), TM::GFP (orange, n=11), and GFP (green, n=7) after photobleaching half of the PCMC. **(f)** Fluorescence recovery of GCY-22::GFP (blue, n=6), TM::GFP (orange, n=7), and GFP (green, n=6) after photobleaching half of the tip compartment. Scale bars represent 1 μm. Colored areas indicate SD.

We photobleached half the PCMC or tip to assess GCY-22::GFP motility *within* these compartments. GCY-22::GFP fully redistributed within the PCMC in 60 seconds (**Fig. 2c**) whereas tip recovery was slower at 3-4 minutes (**Fig. 2d**). These experiments demonstrate lateral diffusion, and no bound fraction, within the PCMC and tip compartments.

To place these results in context, we measured the redistribution of free GFP and of GFP fused to the transmembrane domain of GCY-22 (TM::GFP). After photobleaching half the PCMC, GFP fluorescence recovered quickly (∼90% after 8 seconds). Recovery of TM::GFP was much slower (∼35% after 20 seconds) and comparable to GCY-22::GFP recovery, possibly due to protein crowding within the PCMC (**Fig. 2e**).

After photobleaching half of the cilium, free GFP fluorescence recovered very quicly (∼100% after 2 seconds). TM::GFP recovery was slightly slower (∼80% after 3 seconds), reflecting slower diffusion of transmembrane proteins. In contrast, GCY-22::GFP recovered minimally 10 seconds after photobleaching, indicating the presence of a distinct membrane compartment limiting GCY-22::GFP motility (**Fig. 2f**).

### Ciliary tip compartment of GCY-22::GFP depends on active IFT

Because the proximal region of the axoneme (middle segment) and part of the distal segment seemed void of GCY-22::GFP, we hypothesized that it might be actively transported by IFT. In *C. elegans*, anterograde IFT relies on kinesin-II and OSM-3 (ortholog of mammalian KIF17) in the middle segment and OSM-3 alone in the distal segment^3,4,16,35^.

Using time-lapse microscopy and kymogram analysis, we identified GCY-22::GFP particles moving between the PCMC and ciliary tip (**Fig. 3a**). Two-color imaging of GCY-22::GFP and an mCherry-tagged OSM-3 showed overlapping anterograde tracks (**Fig. 3b**). On average, we observed 37.75 (± 8.23) anterograde OSM-3::mCherry tracks/minute compared to 5.93 anterograde GCY-22::GFP tracks/minute, suggesting that a subset of IFT particles transport GCY-22::GFP (**Table 1, Supplementary Table 1**). Anterograde OSM-3::mCherry particles often move along the full length of the cilium (average track length 2.75 ± 0.54 μm); in contrast, GCY-22::GFP tracks spanned half the cilium or shorter (average length 1.0 ± 0.4 μm, *P* <0.001, **Fig. 3c**). Retrograde GCY-22::GFP particles displayed short tracks and occasionally tracks spanning the entire axoneme (average length 1.3 ±0.8 μm), significantly shorter than retrograde tracks of the tdTomato tagged dynein subunit XBX-1 (average length 2.12 ± 0.59 μm, *P* <0.001) (**Fig. 3c**).

**Table 1.** Quantification of GCY-22::GFP tracks in the cilium in different mutant backgrounds.

| Genotype | # animals with tracks (n) | P-value<sup>a</sup> | Total min. analyzed | Total anterograde tracks (tracks/min) | Total retrograde tracks (tracks/min) |
| --- | --- | --- | --- | --- | --- |
| Wild type | 17(18) | - | 22.75 | 135 (5.93) | 146 (6.42) |
| *kap-1* | 16(18) | 1 | 18 | 179 (9.94) | 127 (7.06) |
| *osm-3* | 7(13) | 0.026 | 19.5 | 33 (1.7) | 18 (0.9) |
| *bbs-8* | 12(17) | 0.094 | 28.5 | 65 (2.3) | 25 (0.9) |
| *daf-25* | 0(18) | <0.001 | - | - | - |
| *mks-5* | 5(14) | <0.001 | 21 | 35 (1.7) | 9 (0.4) |
| *mks-5; daf-25* | 4(21) | <0.001 | 17.5 | 22 (1.3) | 6 (0.3) |
| *gcy-22(RD3>ala)* | 0(16) | <0.001 | - | - | - |
| *rdl-1* | 9(13) | 0.136 | 19.5 | 52 (2.7) | 63 (3.2) |

<sup>a</sup> $\chi^2$ -test was used to compare distributions of animals with and without GCY-22::GFP tracks to WT.

**Figure 3.**
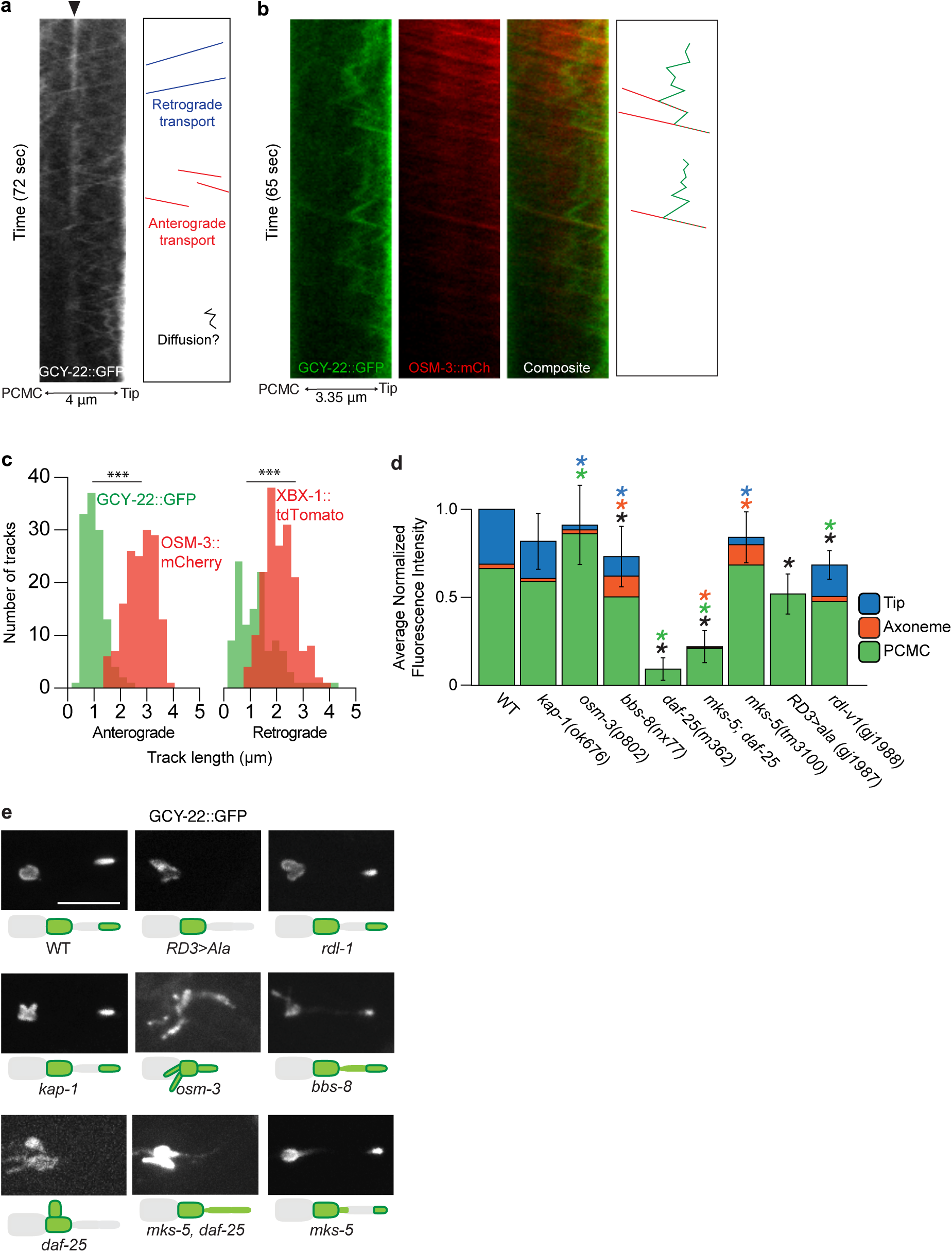
The GCY-22-containing cilium tip compartment is actively maintained by IFT. **(a)** Kymogram of GCY-22::GFP showing anterograde and retrograde ciliary transport and possible diffusion tracks. Black arrowhead indicates stationary signal. **(b)** Kymograms showing partial overlap of IFT tracks of GCY-22::GFP (green) and of the mCherry-tagged anterograde IFT motor protein OSM-3 (red). **(c)** Quantification of track length of GCY-22::GFP (green, n=13 cilia), OSM-3::mCherry (left graph, red, n=7 cilia), and tdTomato-tagged retrograde IFT motor protein XBX-1 (right graph, red, n=24 cilia). Asterisks indicate significant difference in length (*P*-value: * <0.05, ** <0.01, *** <0.001, two-tailed Mann-Whitney U test). **(d)** Quantification of normalized fluorescence intensity in different mutant backgrounds of GCY-22::GFP in the PCMC, axoneme and cilium tip. Distribution of fluorescence between the PCMC (green), axoneme (orange) and cilium tip (blue). Significant differences (P-value <0.05) are indicated by colored asterisks (black is total cilium fluorescence, ANOVA followed by pairwise t-test with Holm correction), n≥5. **(e)** Representative fluorescence images and schematics showing localization of GCY-22::GFP in PCMC and cilium in different mutant backgrounds. Scale bar represents 5 μm.

In many animals, stationary fluorescence signals that indicate paused particles are observed, likely at the transition between the middle and distal segments (**Fig. 3a**). Interestingly, GCY-22::GFP tracks frequently reversed from retrograde to anterograde, docking with an anterograde OSM-3::mCherry track and moving towards the tip (**Fig. 3b**). Notably, this behavior was not observed for IFT proteins. These data suggest that GCY-22::GFP particles moving away from the tip are transported back by anterograde IFT, keeping GCY-22::GFP concentrated at the tip.

To study the contribution of IFT to cilium tip localization, we visualized the localization of GCY-22::GFP in different IFT mutant backgrounds. Disrupting the kinesin-II subunit KAP-1, which plays a role in IFT particle entry into the cilium but does not affect cilium length^3,16,35^, did not influence ciliary GCY-22::GFP levels or localization (**Fig. 3d,e**). In contrast, *osm-3* mutants lacking a distal segment showed increased levels of GCY-22::GFP in the PCMC and along the remaining axoneme but no GCY-22::GFP tip compartment (**Fig. 3d,e**). Disruption of the BBSome subunit BBS-8^36^ reduced overall ciliary levels of GCY-22::GFP, including at the tip, while increasing its localization along the length of the cilium (**Fig. 3d,e**). These changes in GCY-22::GFP localization correlate with a decrease in GCY-22::GFP tracks (**Table 1**). Taken together, our results show that kinesin-II is not required for import of GCY-22::GFP into the cilium, and that both OSM-3-mediated IFT and the BBSome are important for transport into the cilium and enrichment at the tip.

To test if IFT is responsible for the ciliary tip localization of GCY-22::GFP, we used NaN_3_ to stop ATP production, and thus IFT, over time^37^. OSM-3::mCherry particles slowed down and became less frequent after 10 minutes of treatment (**Fig. 4a,b**). IFT capacity, defined as the number of IFT tracks multiplied by speed, reached 0 after 30-50 minutes (**Fig. 4b**). During treatment, cilium length steadily shortened, while the GCY-22::GFP tip compartment extended into the cilium as early as 10 minutes in some animals and completely collapsed after 20-30 minutes (**Fig. 4a,c**). Importantly, loss of cilium tip localization after 30 minutes of NaN_3_ treatment was reversible, with complete recovery within 16 minutes (**Fig. 4d**). These experiments suggest that a minimal IFT capacity is needed to maintain GCY-22::GFP tip localization, although we cannot exclude an IFT-independent mechanism caused by the NaN_3_ treatment.

**Figure 4.**
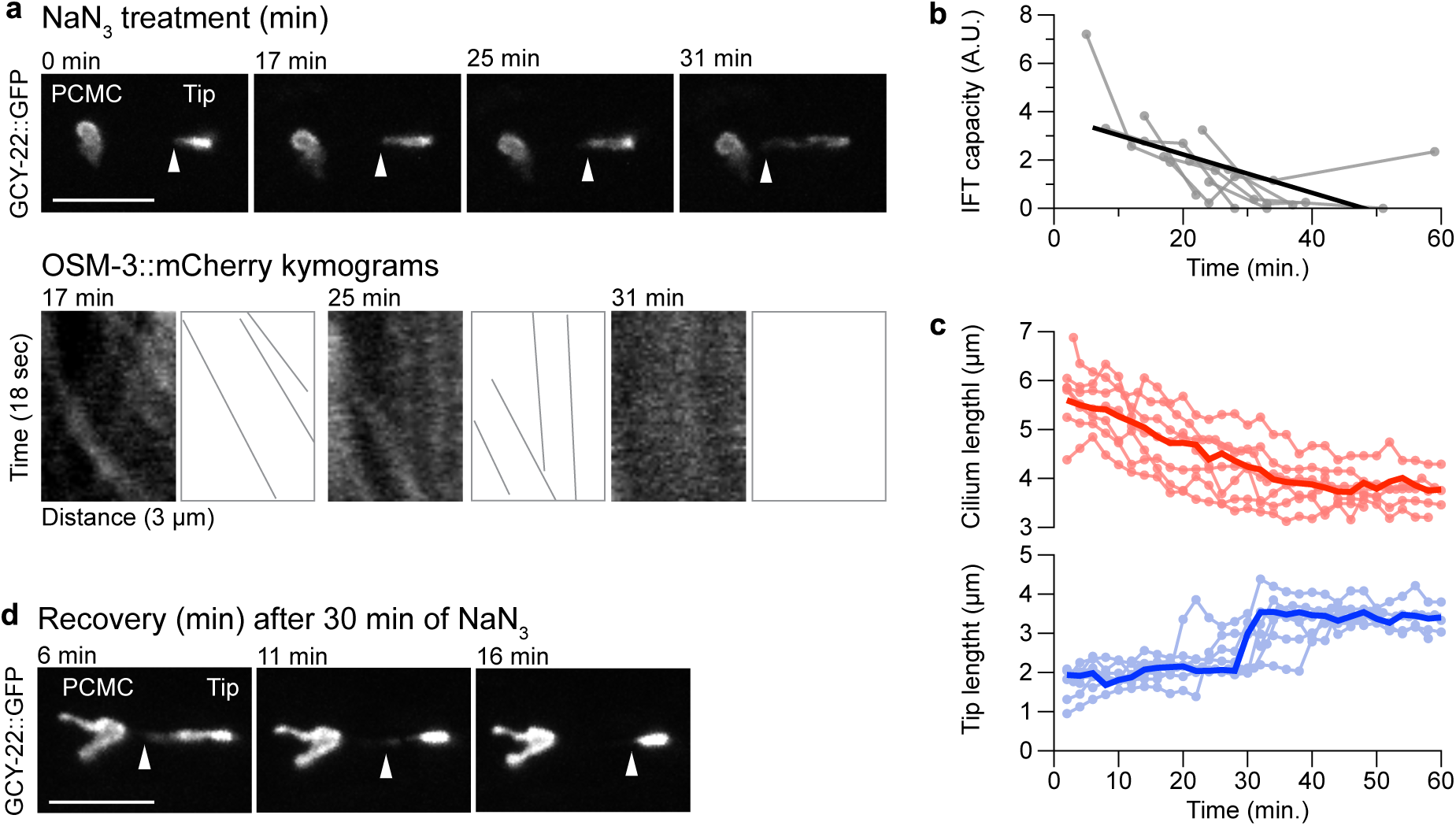
Maintenance of the GCY-22-containing cilium tip compartment is ATP dependent. **(a)** Fluorescence images of GCY-22::GFP showing cilium tip collapse and kymograms of OSM-3::mCherry showing IFT arrest during NaN_3_ treatment. Asterisk indicates PCMC, arrowheads indicate proximal end of tip compartment. **(b)** Quantification of distal segment IFT capacity (number of tracks multiplied by speed) during NaN_3_ treatment (n=7) and linear regression (black line, R^2^=0.37). **(c)** Quantification of cilium length (red) and cilium tip (blue) during NaN_3_ treatment. Darker lines show time-normalized, average result (n=7). **(d)** Representative fluorescence images showing recovery of the cilium tip compartment after 30 min of NaN_3_ treatment (n=5). Arrowheads indicate proximal end of tip compartment. Scale bars represent 5 μm.

Taken together, these experiments show that IFT plays an important role in actively maintaining the high GCY-22 concentration at the ciliary tip.

### GCY-22 trafficking and cilium import is regulated by the transition zone and requires DAF-25

The accumulation of GCY-22::GFP at the PCMC suggests that it is prevented from readily diffusing into the cilium. We therefore tested whether disruption of the TZ influences its ciliary entry. Surprisingly, loss of MKS-5 (mammalian RPGRIP1/RPGRIP1L ortholog), which removes all known proteins and characteristic Y-link structures from the TZ^13,14,38^, has a relatively subtle effect on GCY-22::GFP localization—it mislocalizes at the TZ and base of the axoneme, and displays a reduced amount at the cilium tip (**Fig. 3d,e**). The lack of increased entry suggested a mechanism other than the TZ in regulating GCY-22::GFP ciliary entry.

The Ankmy2 protein DAF-25 is known to be required for ciliary localization of guanylate cyclases^38–40^, cyclic nucleotide gated channels and GPCRs^18,38^, but not for IFT^39^. In *daf-25* mutants, GCY-22::GFP did not enter the cilium, and no IFT tracks of GCY-22::GFP or ciliary tip localization was observed (**Fig. 3d,e, Table 1**). Interestingly, *daf-25* animals showed lower levels of GCY-22::GFP at the PCMC, very few moving vesicles and diffuse fluorescence in the dendrite, and more diffuse localization in the cell body, which did not overlap with the trans-Golgi marker APT-9::mCherry (**Fig. 3e, Supplementary Fig. 1-3**). To test whether DAF-25 functions together with the TZ to traffic GCY-22::GFP into the cilium, we generated a *daf-25; mks-5* mutant strain. In this double mutant, GCY-22::GFP was present at low levels in the cilium but not at the tip (**Fig. 3d,e**). Very few IFT tracks of GCY-22::GFP were visible, potentially explaining the lack of cilium tip localization (**Table 1**). Because OSM-3-mediated IFT is not affected in *daf-25; mks-5* animals (**Supplementary Table 1**), MKS-5 and DAF-25 may be required to link GCY-22::GFP to the IFT machinery, allowing for the formation of the cilium tip compartment.

Together, our data reveals that DAF-25 is required for GCY-22::GFP import across the TZ and into the cilium, and together with the core TZ scaffolding protein MKS-5 is involved in loading GCY-22::GFP as IFT cargo.

### GCY-22::GFP localization requires its dimerization and RD3-associated domains

To identify protein domains required for GCY-22::GFP ciliary trafficking, we generated a series of deletion constructs. Several domains can be recognized in GCY-22 (**Fig. 5**), including an extracellular receptor domain which possibly provides specificity for Cl^-^ions^28,29^. These experiments showed that its dimerization and RD3 domains are required for cilium entry (**Fig. 5**).

**Figure 5.**
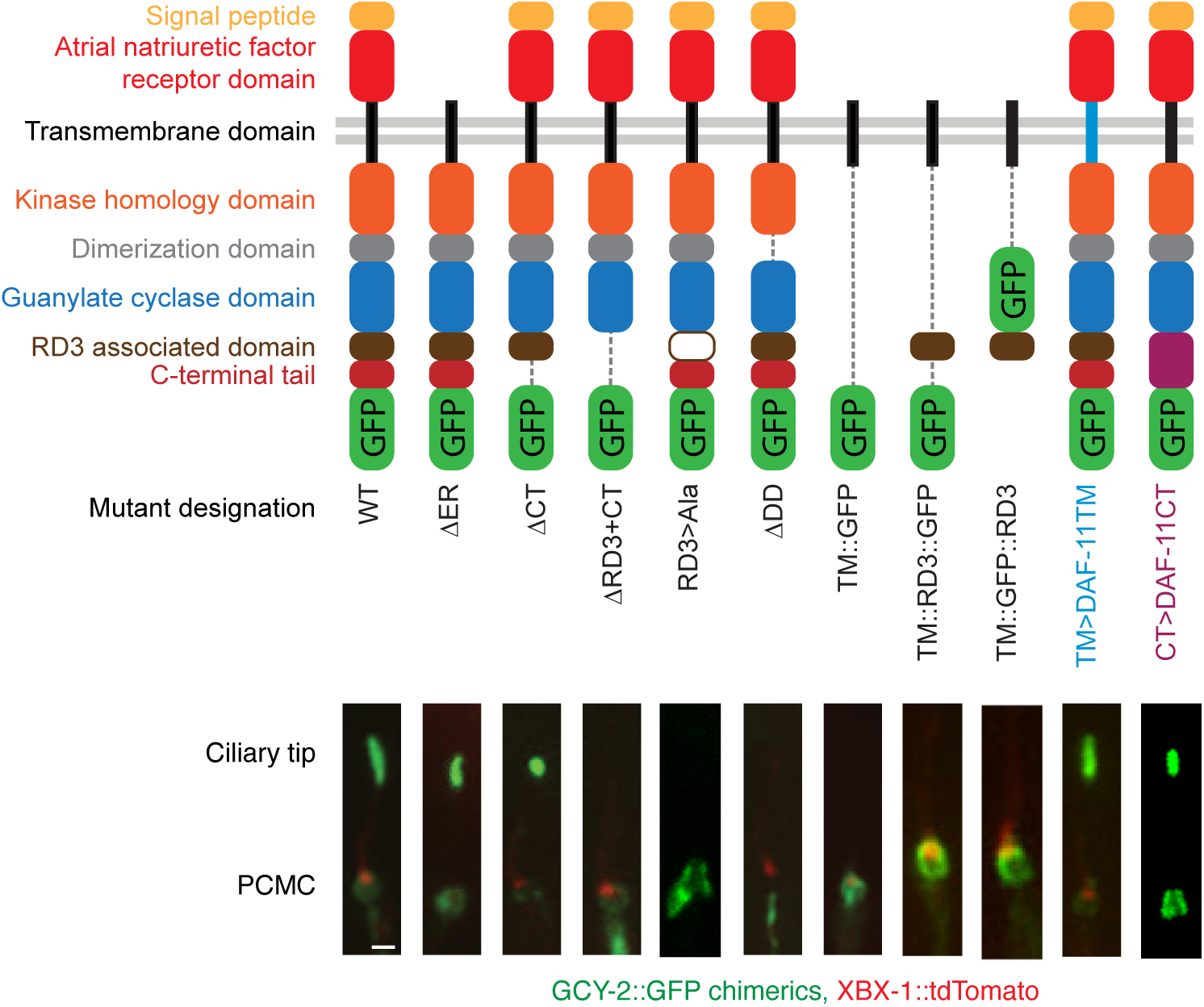
Cilium tip localization of GCY-22::GFP requires the dimerization and RD3-associated domains. Schematic of GCY-22::GFP wild-type and different chimeric proteins showing deleted domains: extracellular receptor domain (ΔER), C-terminal tail (ΔCT), RD3-associated domain (ΔRD3), dimerization domain (ΔDD), transmembrane domain only (TM::GFP), transmembrane domain and RD3-associated domain fused to GFP in two orientations (TM::RD3::GFP and TM::GFP::RD3) and chimeric proteins with GCY-22 domains replaced with corresponding DAF-11 domains, transmembrane domain (TM>DAF-11TM) and CRISPR/Cas9 based C-terminal tail (CT>DAF-11CT). Fluorescence images showing corresponding localization of GCY-22::GFP versions (green) and XBX-1::tdTomato (red).

The RD3 domain shows homology to the RD3-binding domain identified in the mammalian guanylate cyclase GUCY2D/GC1^41,42^. In a *gcy-22 loss-of-function* background, GCY-22(ΔRD3+CT)::GFP localized at the PCMC but did not enter the cilium (**Fig. 5**). To confirm that the RD3 domain is required for cilium import, we replaced 8 residues (W1042-I1049) of this domain with alanines using CRISPR/Cas9, to create GCY-22(RD3>Ala)::GFP. This mutation completely abolished ciliary entry (**Fig. 3d,e**). More diffuse fluorescence in the dendrite and cell body was also observed, reflecting a potential trafficking defect of GCY-22(RD3>Ala)::GFP (**Supplementary Fig. 3**).

Next, we tested if the RD3 domain is *sufficient* for cilium entry by generating two RD3::GFP constructs fused with the GCY-22 transmembrane (TM) domain (TM::GFP::RD3 and TM::RD3::GFP). These proteins showed diffuse localization in the cell body, some dendritic transport, PCMC localization, and a very weak ciliary signal with no evidence of IFT transport (**Fig. 5, Supplementary Fig. 4**). These results suggest that the RD3 domain is not sufficient for correct routing and import in the cilium.

The *C. elegans* RD3 orthologue, RDL-1, influences the trafficking of GCYs to the PCMC and cilium^43^. Mutating *rdl-1* reduced GCY-22::GFP levels in the PCMC but not cilium tip (**Fig. 3d,e**). In addition, GCY-22::GFP was more diffuse in the cell body and dendrite (**Supplementary Fig. 1,3**), in agreement with RDL-1 regulating an early trafficking pathway for GCYs^43^.

In contrast to GCY-22::GFP, the guanylate cyclase DAF-11 localizes along the entire cilium in ASI, ASJ and ASK neurons^39,44^. We swapped the GCY-22 TM domain or C-terminal end, starting at the highly conserved W residue at the start of the RD3 domain, with those of DAF-11 (TM>DAF-11TM and CT>DAF-11CT, respectively). Both chimeric proteins localized to the PCMC and cilium tip, suggesting that the TM domains and C-termini of GCY-22 and DAF-11 do not regulate their sub-ciliary localization (**Fig. 5**). In addition, these results suggest that the C-terminal region of these guanylate cyclases functions cell independently in cilium import.

Together, our data show that GCY-22 ciliary entry and tip localization requires the dimerization and RD3 domains, and likely involve the combined action of more than one domain.

### GCY-22::GFP cilium tip compartment is required for high NaCl sensitivity

ASE neurons express several receptor-type guanylate cyclases, GCY-22 and GCY-14 appearing to be most important for detecting NaCl^26,28,29^. We tested *gcy-22* and *gcy-14* loss-of-function mutants and a *gcy-14; gcy-22* double mutant in a NaCl chemotaxis quadrant assay. Animals are tested for their preference for a particular NaCl concentration versus no NaCl^45,46^. Deleting *gcy-22*, but not *gcy-14*, significantly affected chemotaxis to NaCl (**Fig. 6a**). Chemotaxis by the double mutant was not significantly different from the *gcy-22* single mutant, suggesting that GCY-22 is more important than GCY-14 in our assay (**Fig. 6a**).

**Figure 6.**
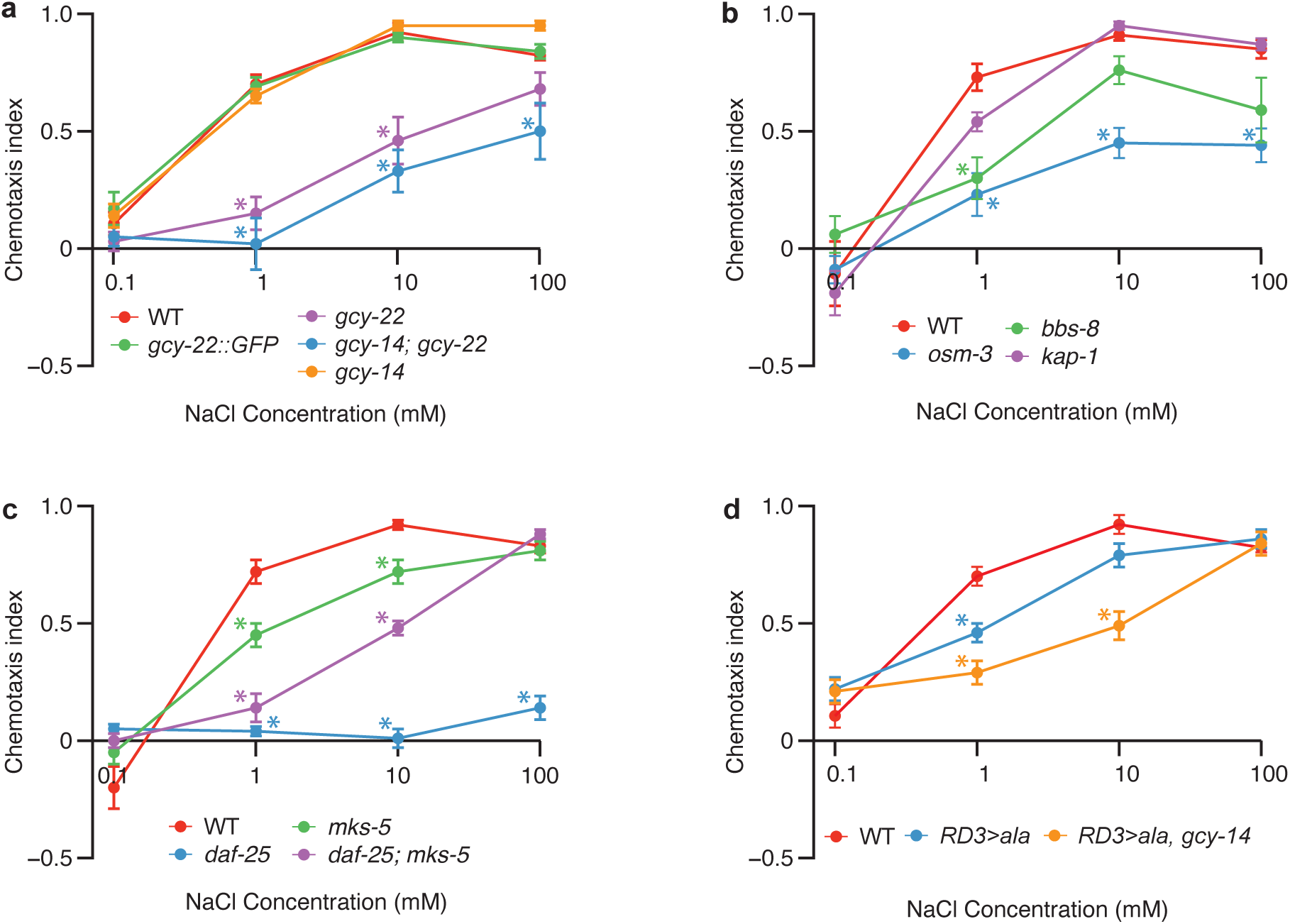
Cilium tip localization of GCY-22 is required for sensitive detection of NaCl. **(a)** Chemotaxis indexes of different mutants showing wild type response of *gcy-22::GFP* animals and involvement of GCY-22. **(b)** Chemotaxis to 1, 10 and 100 mM NaCl requires the IFT anterograde motor OSM-3, and the BBSome subunit BBS-8 for chemotaxis to 1 mM. **(c)** Chemotaxis to 1, 10, and 100 mM NaCl requires the Ankmy2 protein DAF-25, and the TZ component MKS-5 for chemotaxis to 1 and 10 mM. **(d)** Chemotaxis to 1 and 10 mM NaCl requires cilium tip localization of GCY-22 and involves GCY-14. Asterisks indicate significant difference compared to wild type. ANOVA followed by pairwise t-test with Holm correction, n ≥ 6 assays. Full statistical analysis can be found in Supplementary table 2.

To determine if GCY-22 cilium tip localization is important for *C. elegans’* sensitivity to NaCl, we first tested IFT mutants for their chemotaxis response. *kap-1* mutant animals showed a wild-type response to NaCl (**Fig. 6b**), consistent with the normal tip localization of GCY-22::GFP. *bbs-8* mutant animals showed reduced chemotaxis to 1 mM NaCl (P <0.01; **Fig. 6b**), which correlates with the reduced cilium tip levels of GCY-22::GFP. *osm-3* mutant animals, which have short cilia where GCY-22::GFP localizes along its entire length, showed the strongest chemotaxis defect at all NaCl concentrations tested (P<0.01; **Fig. 6b**).

Next, we tested *daf-25* mutant animals, which lack ciliary GCY-22::GFP. These animals showed a strong chemotaxis defect, with only a modest response to 100 mM NaCl (**Fig. 6c**). This suggests that *daf-25* is required for ciliary import of proteins essential for chemotaxis, and consistent with the mislocalization of guanylate cyclases and CNGs^18,39,40^. Surprisingly, disrupting the TZ in the *daf-25* mutant (*daf-25; mks-5* double mutant) resulted in a partially-restored response to 10 mM NaCl, and a wild-type response to 100 mM (**Fig. 6c**). These responses roughly correlate with the levels of ciliary GCY-22::GFP, where the increased amount in the double mutant appears sufficient for detecting high NaCl concentrations. However, the possibility remains that other proteins that play a role in NaCl sensory transduction are also affected.

To specifically test the contribution of GCY-22::GFP ciliary tip localization to NaCl sensation, we tested the RD3>Ala mutant animals for their chemotaxis response. These animals showed a mild chemotaxis defect at 1 mM NaCl (P=0.023), but wild-type responses to 10 and 100 mM NaCl (**Fig. 6d**), suggesting that a small amount of GCY-22 could still be present in the cilium. Additionally, a chemotaxis defect in these animals could be masked by functional redundancy. To test this latter possibility, we made a *gcy-14; gcy-22(RD3*>*Ala)* double mutant and observed a stronger chemotaxis defect at 10 mM NaCl (P=0.020) compared to *gcy-22(RD3*>*Ala)* single mutants (**Fig. 6d**). Interestingly, *gcy-14; gcy-22(RD3*>*Ala)* animals showed stronger chemotaxis to 100 mM NaCl than *gcy-14; gcy-22* animals (P=0.004), indicating that the GCY-22::RD3>Ala protein is functional in detecting NaCl.

Together these experiments show that a high level of GCY-22 is required at the cilium tip for detecting, and efficient chemotaxis to, low NaCl concentrations.

## Discussion

Some signaling proteins localize to specific ciliary subdomains, whereas others distribute along the length of the cilium, suggesting that their specific localization patterns are functionally important. Our understanding of how the signaling protein localization within the cilium is regulated remains limited. Here, we identified mechanisms involved in the localization of the putative Cl^-^ receptor GCY-22 to a unique ciliary tip domain in the ASER neuron of *C. elegans* and provide evidence that this specific localization is essential for its function as a highly sensitive NaCl sensor.

Our results suggest that IFT is the primary driving force behind GCY-22 localization to the cilium tip. First, we identified DAF-25 as essential for ciliary import. DAF-25 functions together with the TZ, by way of MKS-5, to load GCY-22 as cargo onto IFT particles. Second, dual-color imaging showed co-localization of GCY-22::GFP with OSM-3-kinesin in a subset of IFT particles, suggesting association between GCY-22::GFP and certain IFT particles or ‘trains’. Strikingly, many GCY-22::GFP particles moving towards the ciliary base were picked up by anterograde IFT trains and relocalized to the tip. This behavior, not previously observed for other IFT-associated proteins, maintains a high density of receptor molecules within the tip compartment. How this is regulated—for example, whether this depends on differential affinity for anterograde or retrograde IFT machinery—remains to be determined. Finally, interfering with IFT diminishes the accumulation of GCY-22::GFP at the tip. This indicates that IFT plays a crucial role in actively maintaining a specialized signaling compartment at the cilium tip.

Interestingly, our FRAP assays indicate the presence of a distinct tip membrane compartment with properties different from the rest of the cilium. GCY-22 might therefore bind other, yet-to-be-identified protein(s) at the cilium tip. Similarly, how Hedgehog signaling components remain confined to the cilium tip is unknown^47^. However, we found a high density of GCY-22::GFP molecules at the cilium tip, which slows lateral diffusion^48^. This may explain, independent of tethering by other proteins, the slow redistribution of GCY-22::GFP in the tip compartment.

The high concentration of GCY-22 at the ciliary tip suggests that it forms a highly sensitive detection apparatus, reminiscent of the high densities of rhodopsin in mammalian photoreceptor cells, which enable the detection of single photons^34,49^. Indeed, mislocalization of GCY-22::RD3>Ala::GFP affected chemotaxis to 1 mM NaCl most, suggesting that the ciliary tip compartment is required to navigate small differences in NaCl concentrations, likely important for *C. elegans* in its natural habitat.

In conclusion, our study revealed mechanisms for generating and maintaining a specialized ciliary tip domain that is analogous to the cGMP signaling domain of mammalian ciliary photoreceptors, and ciliary tip domain of the Hedgehog signaling cascade. Our findings suggest that such domains may be broadly used as signaling compartments and should be sought and analysed in mammalian cilia in the context of different signaling pathways, human physiology and disease.

## Methods

### Strains and Constructs

Strains were cultured using standard methods^50^. Alleles used in this research were: *kap-1(ok676), osm-3(p802), bbs-8(nx77), daf-25(m362), mks-5(tm3100), gcy-14 (pe1102), gcy-22(tm2364), dhc-1(ie28[dhc-1::degron::GFP]), ieSi57[eft-3p::TIR1::mRuby::unc-54 3’UTR + Cbr-unc-119(+)]*.

The *p*_*rab-3*_::*mCherry::apt-9* construct was (gift from A. Pasparaki) and used to generate *p*_*rab-*_ *3::mCherry::tram-1*. A 1.5 kb genomic DNA fragment containing *tram-1* was amplified with primers #2581 and #2582 and used to replace *apt-9. p*_*gpa-4*_::*gcy-22::GFP* was generated by inserting two PCR fragments of 2.5 kb and 2 kb, amplified with primers #3152 and #3154, and primers #3153 and #3151, together containing the genomic *gcy-22* locus, into pGJ325^46^. To generate the *p*_*U6*_::*osm-3_sgRNA* vector we cloned an *osm-3* guide into the *p*_*U6*_::*unc-119::sgRNA* vector^51^. The *p*_*flp-6*_::*mCherry* construct was generated by inserting a 2 kb *flp-6* promoter sequence, amplified with primers #2867 and #2869, into a pPD95.77 (gift from A. Fire) backbone containing *mCherry*. The *osm-3::mCherry* template construct was generated by inserting *mCherry* and two 1.5 kb homology arms, amplified from genomic DNA using primers #2679 and #2643 and primers #2660 and #2646, into the backbone of *p*_*U6*_::*unc-119::sgRNA*. The *p*_*gcy-5*_::*xbx-1::tdTomato* construct was generated by inserting a 325 bp promoter from *gcy-5*, amplified using primers 355 and 356, and a 2.2 kb genomic fragment of *xbx-1*, amplified using primers 326 and 327, into a pPD95.81 vector containing *tdTomato*. The *p*_*gcy-5*_::*gcy-22::GFP* construct was generated by PCR fusion^52^. First, a 4.6 kb genomic sequence of *gcy-22*, amplified using primers #221 and #248, and a 325 bp genomic fragment of the *gcy-5* promoter, amplified using primers #219 and #222, were fused using primers #443 and #215. Subsequently, this product was fused upstream of the GFP-coding cassette, including the *unc-54* 3’-UTR, from pPD95.67 (gift from A. Fire) using primers #220 and #609. The final fusion product was cloned into pGEMT easy.

The ΔER strain was generated by fusing the 426 bp *gcy-5* promoter, amplified with primers #218 and #458, and a 2.6 kb ΔER fragment, amplified from genomic DNA using primers #457 and #248, using primers #219 and #215. This product was fused to *GFP*::*unc-54-3’-UTR* from pPD95.67 using primers #220 and #609. The ΔCT strain was generated by amplifying from *p*_*gcy-5*_::*gcy-22::GFP*, a *p*_*gcy-5*_::*gcy-22::ΔCT* fragment, using primers M13Fwd and #433, and *GFP* using primers #432 and M13Rev, and fusing these products using primers #220 and #609. The ΔRD3+CT strain was generated by fusing a *gcy-22* fragment upstream of the RD3 domain, amplififed using primers #434 and #435, and a fragment containing GFP, amplified using primers #152 and #434, using primers #220 and #609. The ΔDD strain was generated by amplifying fragments upstream and downstream of the DD from *p*_*gcy-*_ *5::gcy-22::GFP* using primers #470 and #219, and primers #469 and #609, and fusing these products using primers #220 and #609. The *TM*>*DAF-11TM* strain was generated by amplifying *p*_*gcy-5*_::*gcy-22::GFP* fragments up and downstream of the *TM*, using primers #440 and #443, and primers #441 and #215. Primers #441 and #440 contained overlapping regions of the *daf-11 TM*. Primers #443 and #215 were used to fuse these two fragments.

The TM::GFP strain was generated by fusing a *gcy-22 TM* fragment, amplified from ΔER using primers #219 and #454, to *GFP* using primers #220 and #609. The *TM::RD3::GFP* strain was generated by amplifying *a p*_*gcy-5*_::*gcy-22(TM)* fragment, amplified from *p*_*gcy-5*_::*gcy-22(TM)::GFP* using primers #220 and #607. Next, a *gcy-22(RD3)* fragment, amplified from genomic DNA using primers #606 and #607, was fused to *GFP::unc-54-3’-UTR* pPD95.77, using primers #608 and #609. Finally, the *p*_*gcy-5*_::*gcy-22(TM)* and *RD3::GFP::unc-54-3’-UTR* fragments were fused using primers #220 and #609. *TM::GFP::RD3* strain was generated by amplifying a *p*_*gcy-5*_::*gcy-22(TM)::GFP* fragment without *unc-54* 3’-UTR from *p*_*gcy-5*_::*gcy-22(TM)::GFP* using primers #220 and #601, and *a gcy-22(RD3)* fragment from genomic DNA using primers #602 and #603. Subsequently, these two fragments were fused using primers #220 and #604.

PCR fusion products were injected with *p*_*gcy-5*_::*xbx-1::tdTomato*, and pRF4::*rol-6(su1006)*^53^ or *p*_*unc-122*_::*GFP*^*54*^ to generate transgenic strains.

### Microinjections

Microinjections were performed using standard methods^55^.

### CRISPR/Cas9

To generate the *osm-3::mCherry* allele, animals were injected with a mixture containing *p*_*U6*_::*osm-3_sgRNA* (45 ng/μl), *p*_*eft-3*_::*cas9-SV40_NLS::tbb-2* (50 ng/μl), pRF4::*rol-6(su1006)* (50 ng/μl), and a plasmid containing an *mCherry* repair template with 1500 bp homology arms (20 ng/μl). Animals were injected and placed on separate 6 cm NGM plates. Three days later, F1 offspring was picked, allowed to self-reproduce, and screened by PCR. We used a *dpy-10* based co-CRISPR method and a PCR-generated repair template of GFP, amplified with primers #3120 and #3121, with 35 bp homology arms to generate the *gcy-22::GFP(gj1976)* allele. We used ssODN repair template #3393, with 35 bp homology arms, and guide g10, to generate the *gcy-22(RD3*>*Ala)::GFP(gj1987)* allele. The *rdl-1(gj1989)* deletion allele was generated using two guides, g14 and g15, and no template.

We used the CRISPR/Cas9 method as described by Dokshin *et al*.^56^ to generate the *gj2113[gcy-22::CT*>*daf-11CT::GFP]* allele. Briefly, a gBlock (IDT) containing the *daf-11CT* repair template was cloned into a pGEM vector. Subsequently, the repair template was amplified using primers #3404 and #3405 with 35 bp homology arms. The repair template, guide (g10), and pRF-4::*rol-6(su1006)* were injected into GJ3452 according to protocol.

A list of strains, primers, and guides used in this research can be found in Supplementary Table 3.

### Microscopy

Animals were immobilized on a 6% agarose pad, using 0.10 μm polystyrene microspheres (Polybead, Polysciences Inc.) and 10 mM Levamisole (Sigma) as an anesthetic in M9 buffer, unless stated otherwise. Fluorescence images were taken using a spinning disc confocal microscope (Nikon Ti-eclipse) with an EM CCD camera (QuantEM512C, Photometrics) and Metamorph Imaging software, unless stated otherwise. Images were analyzed using FIJI software (version 2.0.0).

For FRAP experiments, a FRAP3D unit (ROPER) was used. Pre-bleach, 10 images were taken at 1 second intervals. Post-bleach the following images were taken: for the entire PCMC or tip 4 images at 15 sec. intervals, 4 images at 60 sec. intervals, and 4 images at 5 min. intervals; for half of the PCMC 120 images at 1 sec. intervals; for half of the cilium tip 10 images at 1 sec. intervals and 18 images at 15 sec. intervals; for GFP 200 images at 50 ms intervals (a subset was plotted in **Fig. 2f**); for TM::GFP in the PCMC 20 images at 1 sec. intervals; for TM::GFP in the cilium tip 33 images at 300 ms intervals. For the time-lapse images, animals were immobilized on a 6% agarose pad and M9 containing 10 mM Levamisole. Images were taken at 300 ms intervals and kymograms were generated using the KymographClear 1.0. ImageJ plugin^57^. Kymograms were analyzed using a custom ImageJ plugin written by I. Smal. Dual color time-lapse images were taken using a DV2 beam splitter (MAG Biosystems).

Quantification of GCY-22::GFP fluorescence in different mutant backgrounds, and of GFP molecules at the cilium tip, was performed on a laser scanning confocal microscope (SP5 AOBS, Leica). A dilution series of purified GFP in PBS was used to generate a calibration curve for the GFP fluorescence. Assuming a confocal volume of 0.1 femtoliter the fluorescence intensity of the calibration curve was converted into number of GFP molecules. The integrated fluorescence intensity of the cilia was measured and converted to number of GFP molecules. The membrane density of GFP molecules in the cilium tip was calculated, assuming a membrane surface of 1.56 μm^2^.

### NaN_3_ treatment

To stop ATP production, NaN_3_ (Sigma) was added to a 6% agarose pad at a concentration of 20 mM. Animals were shortly incubated in a drop of M9 containing 10 mM Levamisole (Sigma) before immobilization. Imaging started 10 min after immobilization. To assess the recovery of the cilium tip, animals were incubated in M9 buffer containing 20 mM NaN_3_ for 30 min and subsequently immobilized for imaging without NaN_3_.

### Auxin inducible degradation

For the auxin inducible degradation of DHC-1::GFP::degron, animals were cultured on NMG plates containing 1 mM IAA (Sigma) for 48 hrs prior to imaging.

### Chemotaxis assays

The quadrant assay used to asses chemotaxis to NaCl was adapted from Wicks *et al*. and Jansen *et al*.^45,46^. Briefly, two diagonally opposite quadrants of a sectional petri dish (Star Dish, Phoenix Biomedical) were filled with 13.5 mL buffered agar (1.7% Agar, 5 mM K_2_HPO_4_/KH_2_PO_4_ pH 6, 1 mM CaCl_2_ and 1 mM MgSO_4_) containing NaCl and two diagonally opposite quadrants with 13.5 mL buffered agar without NaCl. Immediately before the assay, the plastic dividers between the quadrants were covered with a thin layer of agar. Age synchronized *C. elegans* populations were washed 3 times for 5 min with CTX buffer (5 mM K_2_HPO_4_/KH_2_PO_4_ pH 6, 1 mM CaCl_2_ and 1 mM MgSO_4_). Approximately 100 animals were placed in the middle of a sectional dish. After 10 min, animals on each quadrant were counted and a chemotaxis index (CI) was calculated for each plate (CI = (# animals on NaCl – # animals not on NaCl)/ total # animals). To determine the CI of a strain, 2 assays per day were performed on at least 3 days.

### Statistics

Statistical analyses were performed using R software, version 3.6.0. IFT track lengths were compared using a Mann-Whitney U test. Comparisons of the chemotaxis indexes, florescence intensities, and OSM-3::mCherry track counts and speeds were performed with a one-way ANOVA, followed by a pairwise t-test with Holm correction. Distributions of animals with and without GCY-22::GFP IFT-like tracks were compared using a Chi^2^-test.

## Acknowledgements

We thank C.L. van der Burght for discussions on statistical analyses. We thank A. Fire, A. Pasparaki and I. Smal for reagents. Some strains were provided by the CGC, which is funded by NIH Office of Research Infrastructure Programs (P40 OD010440), and the Mitani lab through the National Bio-Resource Project of the MEXT, Japan. This work is part of the research program of the Foundation for Fundamental Research on Matter (FOM), which is financially supported by the Netherlands Organization for Scientific Research (NWO). The studies are also funded by the Canadian Institutes of Health Research (CIHR; grants PJT-156042 and MOP-142243). M.R.L. acknowledges a senior scholar award from Michael Smith Foundation for Health Research (MSFHR).

## Author Contributions

S.N.B., M.R.L., and G.J. conceived the study. S.N.B. and S.R. generated strains, performed experiments and analyzed the data. J.J. and C.L. generated strains, performed experiments and analyzed the data for the construct-based domain deletion experiments. S.R. and G.K. designed and performed the experiments and analysed the data for the GFP quantification experiment. S.R. and S.N.B. designed experiments, S.R. performed the experiments, and S.R., S.N.B., and A.B.H. analysed the data for the FRAP experiments. S.N.B., M.R.L. and G.J. wrote the manuscript.

## Figure Legends

**Supplementary Figure 1**. GCY-22::GFP colocalizes with TRAM-1 and APT-9 positive vesicles in the ASER cell bodies of wild-type animals. (a) Fluorescence images of ASER cell bodies showing GCY-22::GFP (green) and the ER-marker TRAM-1::mCherry (red), and colocalization (composite). (b) Fluorescence images of ASER cell bodies showing GCY-22::GFP (green) and the Golgi-marker APT-9::mCherry (red), and colocalization (composite). (c) Fluorescence images showing more diffuse localization of GCY-22::GFP (green) in *daf-25* and *rdl-1* mutant animals, APT-9::mCherry positive vesicles (red) and no colocalization in *daf-25* animals and partial colocalization *rdl-1* animals (composite). Scale bars represent 5 μm.

**Supplementary Figure 2**. Kymograms of dendritic transport of GCY-22::GFP showing vesicular transport in wild-type animals and reduced transport in different mutant backgrounds. Examples of transport (arrowheads) and stationary signal (asterisks) are indicated. Scale bars represent 9 seconds (vertical) and 2 μm (horizontal).

**Supplementary Figure 3**. GCY-22::GFP localization in the cell body, dendrite and cilium of the ASER neurons in wild-type and different mutant backgrounds.

**Supplementary Figure 4**. Fluorescence images of chimeric proteins containing the transmembrane domain and RD3-associated domain (RD3) and GFP in the cell body, dendrite, and cilium of ASER neurons. Kymograms of the dendrite showing diffuse signal and occasional vesicular transport and of the cilium showing diffuse signal only. Asterisks indicate PCMC, scale bar indicates 5 μm.

**Supplementary Figure 5**. Genomic sequences of alleles generated in this research using CRISPR/Cas9.

