## Supplementary files for "Signaling compartment at the ciliary tip is formed and maintained by intraflagellar transport and functions as sensitive salt detector"

Supplementary Figure 1.

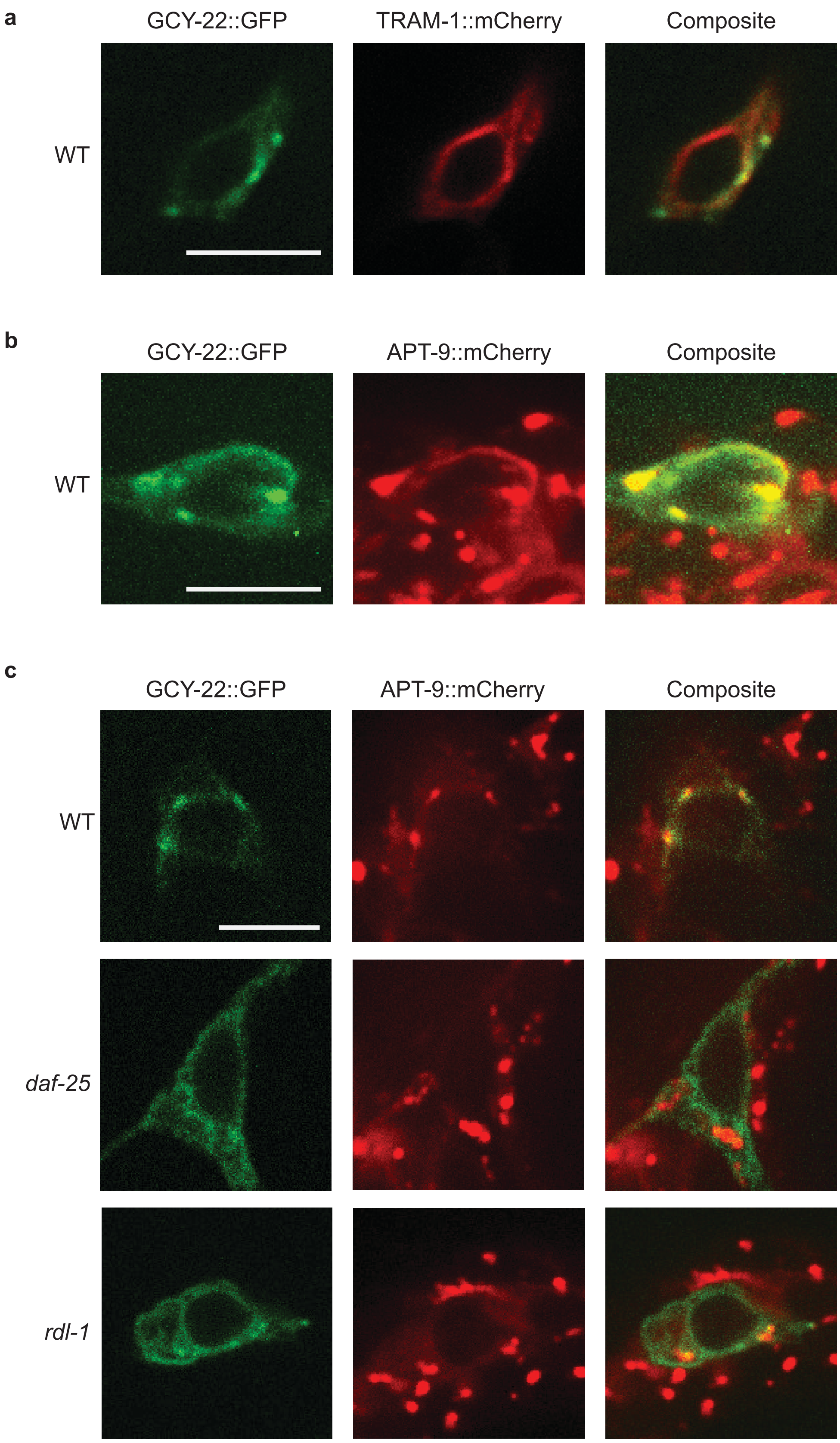

Supplementary Figure 2

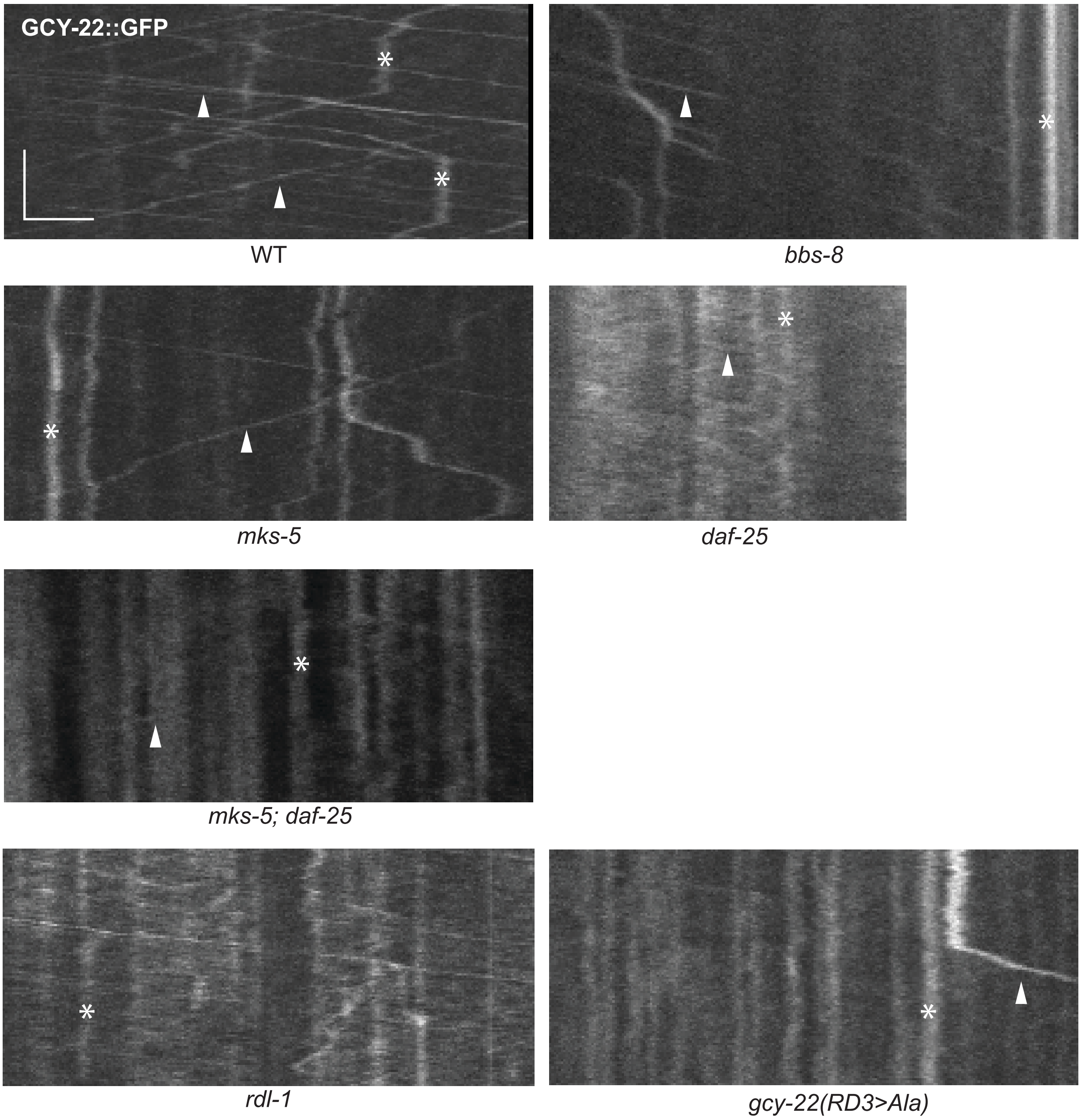

Supplementary Figure 3

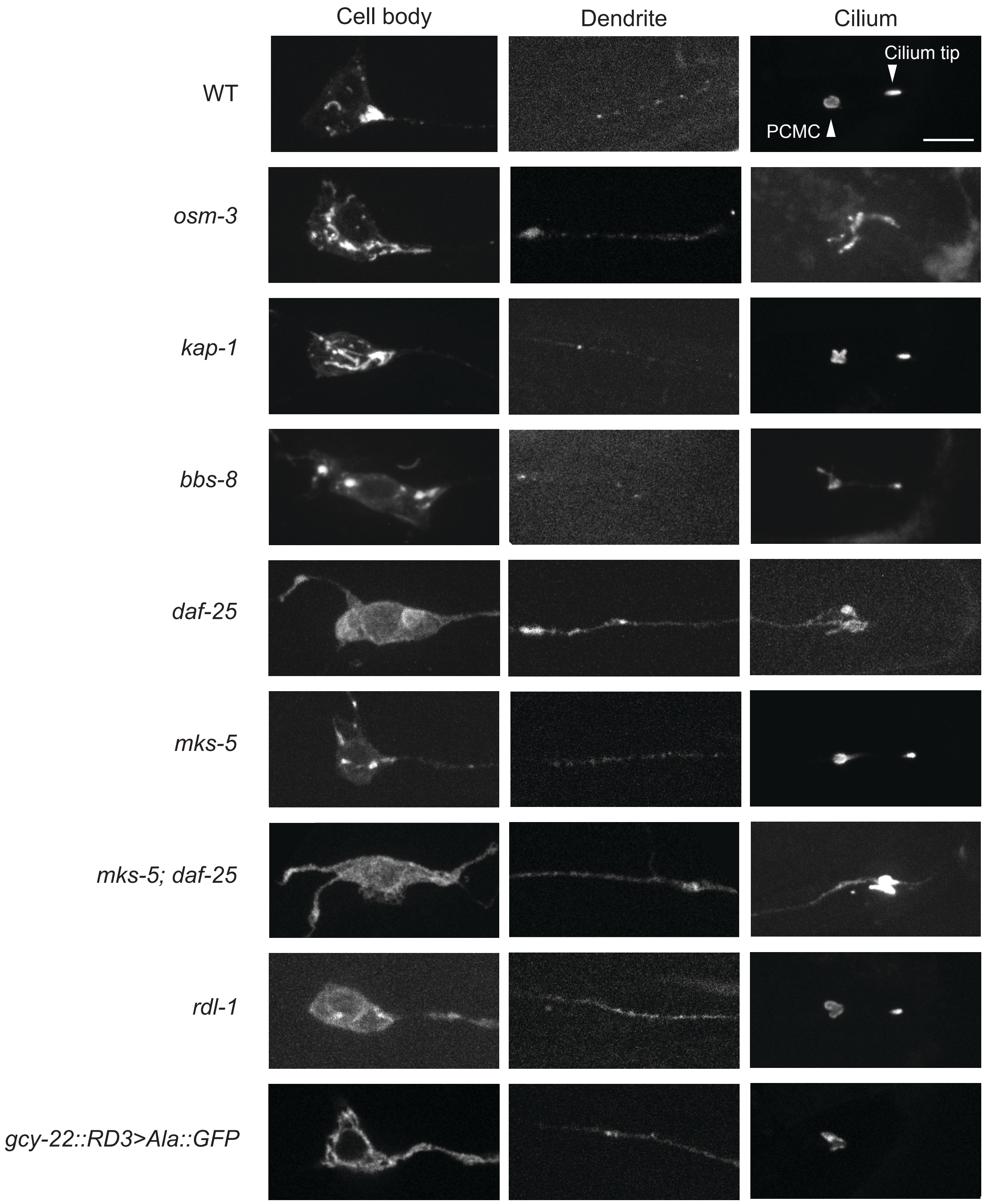

Supplementary Figure 4.

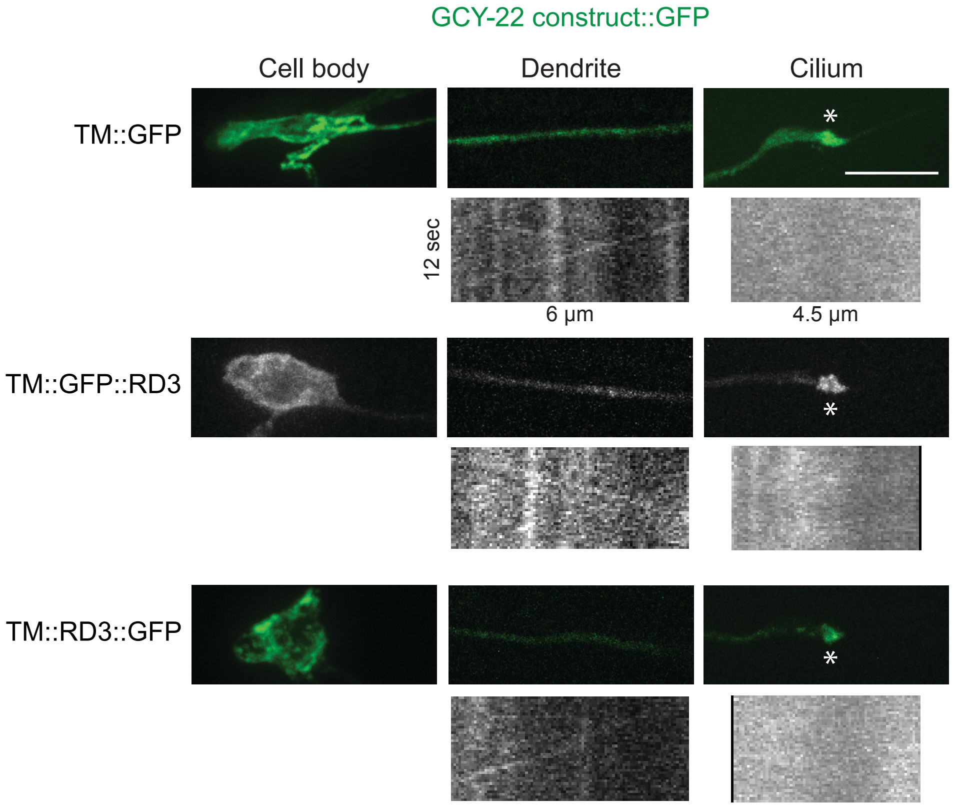

**Supplementary Figure 5. Genomic sequences of alleles generated with CRISPR/Cas9.**

*gcy-22(gj1976[gcy-22::GFP])*

GGAAAAGGTGCAGTTCAAACGCATTGGTTATTGACAGATGATGAAATAGAGGCGAAGGAGAATGGAGAATCTATCGGAGCATCGGGAGCCTCA
GGAGCATCGATGAGTAAAGGAGAAGAATTGTTCACTGGAGTTGTCCCAATCCTCGTCGAGCTCGACGGAGACGTCAACGGACACAAGTTCTCC
GTCTCCGGAGAGGGAGAGGGAGACGCCACCTACGGAAAGCTCACCCTCAAGTTCATCTGCACCACCGGAAAGCTCCCAGTCCCATGGCCAACC
CTCGTCACCACCTTCTGCTACGGAGTCCAATGCTTCTCCCGTTACCCAGACCACATGAAGCGTCACGACTTCTTCAAGTCCGCCATGCCAGAG
GGATACGTCCAAGAGCGTACCATCTTCTTCAAGgtaagtttaaacatatatatactaactactgattatttaaattttcagGACGACGGAAAC
TACAAGACCCGTGCCGAGGTCAAGTTCGAGGGAGACACCCTCGTCAACCGTATCGAGCTCAAGgtaagtttaaacagttcggtactaactaac
catacatatttaaattttcagGGAATCGACTTCAAGGAGGACGGAAACATCCTCGGACACAAGCTCGAGTACAACTACAACTCCCACAACGTC
TACATCATGGCCGACAAGCAAAAGAACGGAATCAAGGTCAACTTCAAGgtaagtttaaacatgattttactaactaactaatctgatttaaat
tttcagATCCGTCACAACATCGAGGACGGATCCGTCCAACTCGCCGACCACTACCAACAAAACACCCCAATCGGAGACGGACCAGTCCTCCTC
CCAGACAACCACTACCTCTCCACCCAATCCGCCCTCTCCAAGGACCCAAACGAGAAGCGTGACCACATGGTCCTCCTCGAGTTCGTCACCGCC
GCCGGAATCACCCACGGAATGGACGAGCTCTACAAGTAA
GCY-22 EXON 17 RD3 associated domain linker GFP intron

*gcy-22(gj1987[gcy-22::RD3>Ala::GFP])*

GGAAAAGGTGCAGTTCAAACGCATGCTGCCGCTGCCGCCGCTGCTGCCGAGGCGAAGGAGAATGGAGAATCTATCGGAGCATCGGGAGCCTCA
GGAGCATCGATGAGTAAAGGAGAAGAATTGTTCACTGGAGTTGTCCCAATCCTCGTCGAGCTCGACGGAGACGTCAACGGACACAAGTTCTCC
GTCTCCGGAGAGGGAGAGGGAGACGCCACCTACGGAAAGCTCACCCTCAAGTTCATCTGCACCACCGGAAAGCTCCCAGTCCCATGGCCAACC
CTCGTCACCACCTTCTGCTACGGAGTCCAATGCTTCTCCCGTTACCCAGACCACATGAAGCGTCACGACTTCTTCAAGTCCGCCATGCCAGAG
GGATACGTCCAAGAGCGTACCATCTTCTTCAAGgtaagtttaaacatatatatactaactactgattatttaaattttcagGACGACGGAAAC
TACAAGACCCGTGCCGAGGTCAAGTTCGAGGGAGACACCCTCGTCAACCGTATCGAGCTCAAGgtaagtttaaacagttcggtactaactaac
catacatatttaaattttcagGGAATCGACTTCAAGGAGGACGGAAACATCCTCGGACACAAGCTCGAGTACAACTACAACTCCCACAACGTC
TACATCATGGCCGACAAGCAAAAGAACGGAATCAAGGTCAACTTCAAGgtaagtttaaacatgattttactaactaactaatctgatttaaat
tttcagATCCGTCACAACATCGAGGACGGATCCGTCCAACTCGCCGACCACTACCAACAAAACACCCCAATCGGAGACGGACCAGTCCTCCTC
CCAGACAACCACTACCTCTCCACCCAATCCGCCCTCTCCAAGGACCCAAACGAGAAGCGTGACCACATGGTCCTCCTCGAGTTCGTCACCGCC
GCCGGAATCACCCACGGAATGGACGAGCTCTACAAGTAA
GCY-22 EXON 17 RD3>Ala linker GFP intron

*gcy-22(gj2113[gcy-22::CT>DAF-11CT])*

GGAAAGGGAGCTGTCCAGACCCACTGGCTGGAAGGAGTAAATGCAGGAGAACAAGTTAAAGTTGTTGAATTCCAAAATGATTTGAACGACGAA
CTTTCAAGAATAATGAAAAAAGATGGAGAGCTTTTAGCTGCAGCAACAGCATTAAAACCAAAAGACAAAATGACGTTGGCGAAAGAAAAAGTG
ATAGCTGAACGAAAAAATGAAGAAGAACGACTCCAACGACAGCAGACATTACAGGAAGCATTGGAAGAACATGAAGAAGAGATAGAAATGAAT
GAAGTACTGGTGGATGAGGATGAAGGTGAAGGAAAACCAAAAGAAGTTGATCTGACAAGTATTGTAAGTACCCAAATGGAAGAGCTAGAAGAT
GAACCAGCAGGTCGAACAATTGGACATGGAAGGCTTGATAGTCAAGCTTCAACTATTCCAGATAACGGAGCAGCTGGAGCCGCTGGAGCAGCC
ATGAGCAAAGGAGAAGAATTGTTCACTGGAGTTGTCCCAATCCTCGTCGAGCTCGACGGAGACGTCAACGGACACAAGTTCTCCGTCTCCGGA
GAGGGAGAGGGAGACGCCACCTACGGAAAGCTCACCCTCAAGTTCATCTGCACCACCGGAAAGCTCCCAGTCCCATGGCCAACCCTCGTCACC
ACCTTCTGCTACGGAGTCCAATGCTTCTCCCGTTACCCAGACCACATGAAGCGTCACGACTTCTTCAAGTCCGCCATGCCAGAGGGATACGTC
CAAGAGCGTACCATCTTCTTCAAGgtaagtttaaacatatatatactaactactgattatttaaattttcagGACGACGGAAACTACAAGACC
CGTGCCGAGGTCAAGTTCGAGGGAGACACCCTCGTCAACCGTATCGAGCTCAAGgtaagtttaaacagttcggtactaactaaccatacatat
ttaaattttcagGGAATCGACTTCAAGGAGGACGGAAACATCCTCGGACACAAGCTCGAGTACAACTACAACTCCCACAACGTCTACATCATG
GCCGACAAGCAAAAGAACGGAATCAAGGTCAACTTCAAGgtaagtttaaacatgattttactaactaactaatctgatttaaattttcagATC
CGTCACAACATCGAGGACGGATCCGTCCAACTCGCCGACCACTACCAACAAAACACCCCAATCGGAGACGGACCAGTCCTCCTCCCAGACAAC
CACTACCTCTCCACCCAATCCGCCCTCTCCAAGGACCCAAACGAGAAGCGTGACCACATGGTCCTCCTCGAGTTCGTCACCGCCGCCGGAATC
ACCCACGGAATGGACGAGCTCTACAAGTAA
GCY-22 EXON 17 SILENT MUTATIONS DAF-11 C-TERMINUS linker GFP intron

*osm-3(gj1932[osm-3::mCherry])*

TATCAACTTCCAAATCGCTATTTCCATCCAAGACCCCAACATTCGATGGTCTTGTCAACGGAGTCGTCTACACTGATGCTCTCTACGAGCGGG
CTCAATCTGCAAAACGACCACCTCGACTAGCTTCTCTGAATCCCAAGGTACCGGTCATGGTGAGCAAGGGCGAGGAGGATAACATGGCCATCA
TCAAGGAGTTCATGCGCTTCAAGGTGCACATGGAGGGCTCCGTGAACGGCCACGAGTTCGAGATCGAGGGCGAGGGCGAGGGCCGCCCCTACG
AGGGCACCCAGACCGCCAAGCTGAAGGTGACCAAGGGTGGCCCCCTGCCCTTCGCCTGGGACATCCTGTCCCCTCAGTTCATGTACGGCTCCA
AGGCCTACGTGAAGCACCCCGCCGACATCCCCGACTACTTGAAGCTGTCCTTCCCCGAGGGCTTCAAGTGGGAGCGCGTGATGAACTTCGAGG
ACGGCGGCGTGGTGACCGTGACCCAGGACTCCTCCCTGCAGGACGGCGAGTTCATCTACAAGGTGAAGCTGCGCGGCACCAACTTCCCCTCCG
ACGGCCCCGTAATGCAGAAGAAGACCATGGGCTGGGAGGCCTCCTCCGAGCGGATGTACCCCGAGGACGGCGCCCTGAAGGGCGAGATCAAGC
AGAGGCTGAAGCTGAAGGACGGCGGCCACTACGACGCTGAGGTCAAGACCACCTACAAGGCCAAGAAGCCCGTGCAGCTGCCCGGCGCCTACA
ACGTCAACATCAAGTTGGACATCACCTCCCACAACGAGGACTACACCATCGTGGAACAGTACGAACGCGCCGAGGGCCGCCACTCCACCGGCG
GCATGGACGAGCTGTACAAGTAA

OSM-3 EXON 10 mCherry

*rdl-1(gj1989)*

GTACCGTATTTTTGGTGGAAAATCGGAAGATATCGCGTCTGGGTAATATTTATAGTTTATCTTTTCACTGAGAATTA---
CTTATTTTTCTAAATTACTTTTAATTTTTATTCAATTTTTGGATTATAAATATTTGTTCTTCACCAACTCCGGCTCGTTTT

38-144 bp upstream of ATG 1685 bp deletion 18-98 bp downstream of STOP

**Supplementary Table 1.** Tracks/min and speed of OSM-3::mCherry IFT tracks in different mutant backgrounds.

| **Genotype** | **Average tracks/min. ±SD** | ***P*-value[a]** | **n** | **Average speed ±SD (μm/sec.)** | ***P*-value[a]** | **n** |
|---|---|---|---|---|---|---|
| Wild type | 37.75 ±8.23 | - | 16 | 0.81 ±0.17 | - | 16 |
| *kap-1(ok676* | 37.08 ±5.07 | 1.000 | 13 | 1.21 ±0.26 | <0.001 | 21 |
| *daf-25(m362)* | 34.25 ±5.26 | 1.000 | 8 | 0.84 ±0.13 | 1.000 | 10 |
| *mks-5(tm3100)* | 30.17 ±5.96 | 0.095 | 8 | 0.73 ±0.13 | 0.626 | 15 |
| *mks-5(tm3100); daf-25(m362)* | 29.83 ±6.67 | 0.078 | 8 | 0.79 ±0.22 | 0.675 | 12 |

[a] Differences compared to wild-type, *P*-values were determined by ANOVA and pairwise t-test with Holm correction.

**Supplementary Table 2.** P-values of pairwise t-tests with Holm correction of chemotaxis assays at different concentrations of NaCl.

| **1 mM NaCl** | WT | *gcy-22::GFP* | *gcy-14(pe1102)* | *gcy-22(tm2364)* | *gcy-14(pe1102); gcy-22(tm2364)* | *gcy-22(RD3>Ala)* |
|---|---|---|---|---|---|---|
| *gcy-22::GFP* | 1.000 | - | - | - | - | - |
| *gcy-14(pe1102)* | 1.000 | 1.000 | - | - | - | - |
| *gcy-22(tm2364)* | 0.000 | 0.000 | 0.000 | - | - | - |
| *gcy-14(pe1102); gcy-22(tm2364)* | 0.000 | 0.000 | 0.000 | 0.616 | - | - |
| *gcy-22(RD3>Ala)* | 0.023 | 0.026 | 0.200 | 0.002 | 0.000 | - |
| *gcy-14(pe1102); gcy-22(RD3>Ala)* | 0.000 | 0.000 | 0.004 | 0.616 | 0.062 | 0.283 |

| **10 mM NaCl** | WT | *gcy-22::GFP* | *gcy-14(pe1102)* | *gcy-22(tm2364)* | *gcy-14(pe1102); gcy-22(tm2364)* | *gcy-22(RD3>Ala)* |
|---|---|---|---|---|---|---|
| *gcy-22::GFP* | 1.000 | - | - | - | - | - |
| *gcy-14(pe1102)* | 1.000 | 1.000 | - | - | - | - |
| *gcy-22(tm2364)* | 0.000 | 0.000 | 0.000 | - | - | - |
| *gcy-14(pe1102); gcy-22(tm2364)* | 0.000 | 0.000 | 0.000 | 0.980 | - | - |
| *gcy-22(RD3>Ala)* | 0.977 | 0.980 | 0.955 | 0.001 | 0.000 | - |
| *gcy-14(pe1102); gcy-22(RD3>Ala)* | 0.000 | 0.001 | 0.001 | 1.000 | 0.955 | 0.020 |

| **100 mM NaCl** | WT | *gcy-22::GFP* | *gcy-14(pe1102)* | *gcy-22(tm2364)* | *gcy-14(pe1102); gcy-22(tm2364)* | *gcy-22(RD3>Ala)* |
|---|---|---|---|---|---|---|
| *gcy-22::GFP* | 1.000 | - | - | - | - | - |
| *gcy-14(pe1102)* | 1.000 | 1.000 | - | - | - | - |
| *gcy-22(tm2364)* | 0.313 | 0.342 | 0.024 | - | - | - |
| *gcy-14(pe1102); gcy-22(tm2364)* | 0.001 | 0.001 | 0.000 | 0.342 | - | - |
| *gcy-22(RD3>Ala)* | 1.000 | 1.000 | 1.000 | 0.168 | 0.000 | - |
| *gcy-14(pe1102); gcy-22(RD3>Ala)* | 1.000 | 1.000 | 1.000 | 0.474 | 0.004 | 1.000 |

| **1 mM NaCl** | WT | *kap-1(ok676)* | *osm-3(p802)* |
|---|---|---|---|
| *kap-1(ok676)* | 0.178 | - | - |
| *osm-3(p802)* | 0.001 | 0.026 | - |
| *bbs-8(nx77)* | 0.002 | 0.077 | 0.534 |

| **10 mM NaCl** | WT | *kap-1(ok676)* | *osm-3(p802)* |
|---|---|---|---|
| *kap-1(ok676)* | 0.543 | - | - |
| *osm-3(p802)* | 0.000 | 0.000 | - |
| *bbs-8(nx77)* | 0.064 | 0.025 | 0.000 |

| **100 mM NaCl** | WT | *kap-1(ok676)* | *osm-3(p802)* |
|---|---|---|---|
| *kap-1(ok676)* | 0.87 | - | - |
| *osm-3(p802)* | 0.01 | 0.01 | - |
| *bbs-8(nx77)* | 0.15 | 0.15 | 0.40 |

| **0.1 mM NaCl** | WT | *daf-25(m362)* | *daf-25(m362); mks-5(tm3100); gcy-22::GFP* |
|---|---|---|---|
| *daf-25(m362)* | 0.037 | - | - |
| *daf-25(m362); mks-5(tm3100); gcy-22::GFP* | 0.106 | 0.944 | - |
| *mks-5(tm3100)* | 0.323 | 0.553 | 0.944 |

| **1 mM NaCl** | WT | *daf-25(m362)* | *daf-25(m362); mks-5(tm3100); gcy-22::GFP* |
|---|---|---|---|
| *daf-25(m362)* | 0.000 | - | - |
| *daf-25(m362); mks-5(tm3100); gcy-22::GFP* | 0.000 | 0.127 | - |
| *mks-5(tm3100)* | 0.004 | 0.000 | 0.001 |

| **10 mM NaCl** | WT | *daf-25(m362)* | *daf-25(m362); mks-5(tm3100); gcy-22::GFP* |
|---|---|---|---|
| *daf-25(m362)* | 0.000 | - | - |
| *daf-25(m362); mks-5(tm3100); gcy-22::GFP* | 0.000 | 0.000 | - |
| *mks-5(tm3100)* | 0.001 | 0.000 | 0.000 |
| **100 mM NaCl** | WT | *daf-25(m362)* | *daf-25(m362); mks-5(tm3100); gcy-22::GFP* |
| *daf-25(m362)* | 0.000 | - | - |
| *daf-25(m362); mks-5(tm3100); gcy-22::GFP* | 0.648 | 0.000 | - |
| *mks-5(tm3100)* | 0.551 | 0.000 | 0.407 |

**Supplementary Table 3.** Strains, guides, and primers used in this research.

| Strain ID | Genotype | Reference |
|---|---|---|
| GJ3452 | *gcy-22(gj1976[gcy-22::GFP])V* | This research |
| GJ3453 | *gcy-22(gj1976[gcy-22::GFP])V; gjEx1980[*$p_{flp-6}$*::mCherry* $p_{elt-2}$*::mCherry]* | This research |
| GJ3470 | *gjEx1986[*$p_{gpa-4}$*::gcy-22::GFP,* $p_{elt-2}$*::mCherry]* | This research |
| GJ3283 | *osm-3(gj1932[osm-3::mCherry])IV* | This research |
| GJ3455 | *osm-3(gj1932[osm-3::mCherry])IV; gcy-22(gj1976[gcy-22::GFP])V* | This research |
| GJ3718 | *dhc-1(ie28[dhc-1::degron::GFP])I; ieSi57[eft-3p::TIR1::mRuby::unc-54 3'UTR + Cbr-unc-119(+)]II; gcy-22(gj1976[gcy-22::GFP])V* | This research |
| GJ3456 | *kap-1(ok676)III; gcy-22(gj1976[gcy-22::GFP])V* | This research |
| GJ3461 | *osm-3(p802)IV; gcy-22(gj1976[gcy-22::GFP])V* | This research |
| GJ3457 | *bbs-8(nx77)V, gcy-22(gj1976[gcy-22::GFP])V* | This research |
| GJ3454 | *daf-25(m326)I; gcy-22(gj1976[gcy-22::GFP])V* | This research |
| GJ3479 | *mks-5(tm3100)II; gcy-22(gj1976[gcy-22::GFP])V* | This research |
| GJ3706 | *daf-25(m326)I; mks-5(tm3100)II; gcy-22(gj1976[gcy-22::GFP])V* | This research |
| GJ3467 | *rdl-1(gj1989)IV; gcy-22(gj1976[gcy-22::GFP])V* | This research |
| GJ3464 | *gcy-22(gj1987[gcy-22::RD3>Ala::GFP])V* | This research |
| GJ3724 | *gcy-22(gj2113[gcy-22::daf-11CT::GFP])V* | This research |
| GJ3723 | *daf-25(gj2112)I; osm-3(gj1932[osm-3::mCherry])IV; gcy-22(gj1976[gcy-22::GFP])V* | This research |
| GJ3814 | *gcy-14(pe1102)V, gcy-22(gj1987[gcy-22::RD3>Ala::GFP])V* | This research |
| GJ2253 | *gcy-14(pe1102)V, gcy-22(tm2365)V* | This research |
| MX1845 | *nxEx933[*$p_{gcy-5}$*::gcy-22::GFP,* $p_{gcy-5}$*::xbx-1::tdTomato, coel::GFP]* | This research |
| MX2071 | *nxEx1063[*$p_{gcy-5}$*::gcy-22(ΔER)::GFP,* $p_{gcy-5}$*::xbx-1::tdTomato, coel::GFP]* | This research |
| MX2077 | *nxEx1069[*$p_{gcy-5}$*::gcy-22(ΔCT)::GFP,* $p_{gcy-5}$*::xbx-1::tdTomato, coel::GFP]* | This research |
| MX2076 | *nxEx1068[*$p_{gcy-5}$*::gcy-22(ΔRD3+CT)::GFP,* $p_{gcy-5}$*::xbx-1::tdTomato, coel::GFP]* | This research |
| MX2208 | *nxEx1143[*$p_{gcy-5}$*::gcy-22(ΔDD)::GFP,* $p_{gcy-5}$*::xbx-1::tdTomato, coel::GFP]* | This research |
| MX2883 | *nxEx2883[*$p_{gcy-5}$*::gcy-22(TM)::GFP,* $p_{gcy-5}$*::xbx-1::tdTomato, pRF4::rol-6(su1006)]* | This research |
| MX2917 | *nxEx2917[*$p_{gcy-5}$*::gcy-22(TM)::gcy-22(RD3)::GFP;* $p_{gcy-5}$*::xbx-1::tdTomato, pRF4::rol-6(su1006)]* | This research |
| MX2915 | *nxEx2915[*$p_{gcy-5}$*::gcy-22(TM)::GFP::gcy-22(RD3);* $p_{gcy-5}$*::xbx-1::tdTomato, pRF4::rol-6(su1006)]* | This research |
| MX2884 | *nxEx2884[*$p_{gcy-5}$*::gcy-22(TM>daf-11TM)::GFP;* $p_{gcy-5}$*::xbx-1::tdTomato, pRF4::rol-6(su1006)]* | This research |
| OH4839 | *gcy-22(tm2364)V* | Ortiz *et al.* 2009 |
| JN1194 | *gcy-14(pe1102)V* | Ortiz *et al.* 2009 |
| MX52 | *bbs-8(nx77)V* | Blacque *et al.* 2004 |
| PR802 | *osm-3(p802)IV* | Snow *et al.* 2004 |
| RB849 | *kap-1(ok676)III* | The *C. elegans* deletion mutant consortium *et al.* 2012 |
| MX111 | *mks-5(tm3100)II* | Huang *et al.* 2011 |
| MX1370 | *daf-25(m326)I* | Jensen *et al.* 2010 |

| Primer # | Use case | Sequence |
|---|---|---|
| 3120 | *gcy-22::GFP* repair template | ATGAAATAGAGGCGAAGGAGAATGGAGAATCTATCGGAGCATCGGGAGCCTCAGG |
| 3121 | *gcy-22::GFP* repair template | ATGAAAGAATTGGATAAATCACTAATAATGTGTTACTTGTAGAGCTCGTCCATTC |
| 3393 | *gcy-22::RD3>Ala* repair template | GATGCTCCGATAGATTCTCCATTCTCCTTCGCCTCGGCAGCAGCGGCGGCAGCGGCAGCATGCGT TTGAACTGCACCTTTTCCCTGAAAAAAAA |
| 2581 | *tram-1* locus | CGCGGACCGGTACATATGGTTAAGCCGCAAGGAGGGTCG |
| 2582 | *tram-1* locus | GATTCTCGAGTTCTTAATTTTTCTTCTTCGAATCGC |
| 3152 | *gcy-22* locus | GTGAACACTTTTCAACAAAGCTTGATGAGTTTCATATCAAAATGTTTTATTTG |
| 3154 | *gcy-22* locus | CTAAACTGGAGAAGTGAACCTGCC |
| 3153 | *gcy-22* locus | GGCAGGTTCACTTCTCCAGTTTAG |
| 3151 | *gcy-22* locus | TCCTTTACTCATTTTTTCTACCGGCGATGCTCCGATAGATTCTCCATTCTCCTTCGC |
| 2867 | *flp-6* promoter | CAACTTGGAAATGAAATAAGCTTGCATGCCCTAGTTCTGGGATTTTGCAAG |
| 2869 | *flp-6* promoter | AAAGTTCTTCTCCTTTACTCATTTTTTCTACCGGTATTCTGGAATAATCATATTG |

| | | |
|---|---|---|
| 2679 | *osm-3* homology arm | CTTGCTCACCATGACCGGTACCTTGGGATTCAGAGAAGCTAG |
| 2643 | *osm-3* homology arm | TTGTAAAACGACGGCCAGTGAATGGAAGTGCTAGCCTAGG |
| 2660 | *osm-3* homology arm | CGAGCTGTACAAGTAAGAATTCTTCGTGTGTACATTGTGATG |
| 2646 | *osm-3* homology arm | GACCATGATTACGCCAAGCTCATATGGAGGCGGTTGGTCTTATTAC |
| 3404 | *daf-11CT* repair template | GAATGCATGAAGTTCCTAAC |
| 3405 | *daf-11CT* repair template | GAGCTTTCCGTAGGTGGCG |
| 355 | *gcy-5* promoter | GGGGACAAGTTTGTACAAAAAAGCAGGCTTTCACCAACTTCCGTGAATCCAAG |
| 356 | *gcy-5* promoter | GGGGACCACTTTGTACAAGAAAGCTGGGTTTTTCATCAGAATAAGTA |
| 326 | *xbx-1* locus | GGGGTACCGATGAACATTTGGGATCTTGC |
| 327 | *xbx-1* locus | GGGGTACCCCTCGAACATTAATTTTTGCG |
| 221 | *gcy-22* locus | AATTACTTATTCTGATGAAAAATGAGTTTCATATCAAAATG |
| 222 | *gcy-5* promoter | CATTTTGATATGAAACTCATTTTTCATCAGAATAAGTAATT |
| 248 | *gcy-22* locus | GATAGATTCTCCATTCTCCTTC |
| 219 | *gcy-5* promoter | GTCGTAGTTGTATTTTATGGAAAGTC |
| 443 | Fusion PCR | CATTTTTGCAATTTCACCAACTTC |
| 215 | Fusion PCR | AGTCGACCTGCAGGCATGCAAGCTTGATAGATTCTCCATTCTCCTTC |
| 220 | Fusion PCR | CACCAACTTCCGTGAATCCAAG |
| 218 | *gcy-5* promoter | GTCGTAGTTGTATTTTATGGAAAGTC |
| 458 | *gcy-5* promoter | CATTTTTCATCAGAATAAGTAATTTTTCG |
| 457 | *gcy-22* ΔER fragment | AATTACTTATTCTGATGAAAAATGCAGATAACAGAAGATCTAG |
| 433 | *gcy-22* ΔCT fragment | AGTCGACCTGCAGGCATGCAAGCTTTATTTCATCATCTGTCAATAAC |
| 432 | GFP | GTTATTGACAGATGATGAAATAAAGCTTGCATGCCTGCAGGTCGACT |
| 434 | *gcy-22* ΔRD3+CT fragment | GTGCAGTTCAAACGCATTGGAAGCTTGCATGCCTGCAGGTCGACT |
| 435 | *gcy-22* ΔRD3+CT fragment | AGTCGACCTGCAGGCATGCAAGCTTCCAATGCGTTTGAACTGCAC |
| 470 | *gcy-22* ΔDD fragment | CAATTTTTCAGCCACTTGCTCCAAATTAGATGCATATTG |
| 469 | *gcy-22* ΔDD fragment | CAATATGCATCTAATTTGGAGCAAGTGGCTGAAAAATTG |
| 454 | *gcy-22* TM fragment | AGTCGACCTGCAGGCATGCAAGCTTTTCTGTGTCTTTCTTTGAATTGATTG |
| 440 | *TM>DAF-11TM* fragment | CATCAATAGAACACCCATAATAACAGCAATTGCAATGATCACATGGTCAGTGAAAGATTTTGGAC |
| 441 | *TM>DAF-11TM* fragment | GTGTTCTATTGATGTTTATTATTATATTAACAACAATACGGAAGTGTGTATTTTTAGTAAGGTATGTATAAC |
| 601 | *TM::GFP::RD3* | TTTGTATAGTTCATCCATGCCATGTG |
| 602 | *TM::GFP::RD3* | CACATGGCATGGATGAACTATACAAAATAGGTGGTTATCAAACAGAGCCA |
| 603 | *TM::GFP::RD3* | TTATTTGCATAGACGCACCCATCG |
| 604 | *TM::GFP::RD3* | GTGTACACAGAAAGGATAAGAGCC |
| 605 | *TM::RD3::GFP* | TTCTGTGTCTTTCTTTGAATTGATTGTC |
| 606 | *TM::RD3::GFP* | CTGGAACGAAAATCCTGATCAACG |
| 607 | *TM::RD3::GFP* | AGTCGACCTGCAGGCATGCAAGCTTGATAGATTCTCCATTCTCCTTCGC |
| 608 | *TM::RD3::GFP* | GACAATCAATTCAAAGAAAGACACAGAAATAGGTGGTTATCAAACAGAGCCA |
| 609 | Fusion to *unc-54* 3'-UTR | GGAAACAGTTATGTTTGGTATATTGGG |

| **guide #** | **Allele** | **Sequence** |
|---|---|---|
| g31 | *gcy-22(gj1976)* | TTGGATAAATCACTAATAAT |
| g10 | *gcy-22(gj1987), gcy-22(gj2113)* | AAGGTGCAGTTCAAACGCAT |
| g15 | *rdl-1(gj1989)* | GGAAAATTGACTGGTTTAGC |
| g14 | *rdl-1(gj1989)* | TTAGAAAAATAAGATATTGT |
| g46 | *daf-25(gj2112)* | TAATGTCGTGGCGAGATCCA |
| g47 | *daf-25(gj2112)* | GTATATTGAGATCGAAATAT |
| g50 | *osm-3(gj1932)* | GAATTATTTGGGATTCAGAG |
